# Coronavirus NSP14 Drives Internal m^7^G Modification to Rewire Host Splicing and Promote Viral Replication

**DOI:** 10.1101/2025.09.03.673802

**Authors:** Ensueño Esmeralda Sáenz Altamirano, Chien-Hsin Huang, Yueh-Lin Tsai, Tiffany J. Tzeng, Natália Fagundes Borges Teruel, Isha Pandey, Neelam Oswal, Y. Grace Chen, Ching-Wen Chang, Ricardo Rajsbaum, Lok-Yin Roy Wong, Ivan R. Corrêa, Jack Chun-Chieh Hsu

**Affiliations:** Center for Virus-Host-Innate-Immunity, Institute for Infectious and Inflammatory Diseases, New Jersey Medical School, Rutgers, The State University of New Jersey, Newark, NJ 07103, USA; New England Biolabs Inc., Beverly, MA 01915, USA; Department of Immunobiology, Yale University School of Medicine, New Haven, CT 06520, USA; Center for Discovery and Innovation, Hackensack Meridian Health, Nutley, NJ 07110, USA; Department of Medicine, New Jersey Medical School, Rutgers New Jersey Medical School, Rutgers, The State University of New Jersey, Newark, NJ 07103, USA; Department of Microbiology, Biochemistry and Molecular Genetics, Rutgers New Jersey Medical School, Rutgers, The State University of New Jersey, Newark, NJ 07103, USA

## Abstract

SARS-CoV-2, the causative agent of COVID-19, manipulates host gene expression through multiple mechanisms, including disruption of RNA processing. Here, we identify a novel function of the viral nonstructural protein 14 (NSP14) in inducing *N*7-methylguanosine (m^7^G) modification in the internal sequences of host mRNA. We demonstrate that NSP14 catalyzes the conversion of guanosine triphosphate (GTP) to m^7^GTP, which is subsequently incorporated into mRNA by RNA polymerase II, resulting in widespread internal m^7^G modification. This activity is dependent on NSP14’s *N*7-methyltransferase (*N*7-MTase) domain and is enhanced by interaction with NSP10. Internal m^7^G modification by NSP14 predominantly occurs in the nucleus and is conserved across alpha-, beta- and gamma-coronaviruses. Mechanistically, we show that this RNA modification disrupts normal splicing by promoting intron retention and generating novel splice junctions, particularly in genes regulating genome stability, RNA metabolism and nuclear processes. Importantly, inhibition of m^7^G modification, through pharmacological targeting of NSP14 or RNA polymerase II, impairs SARS-CoV-2 replication, indicating that the virus hijacks host transcriptomic machinery to support infection. Our findings reveal a previously unrecognized epitranscriptomic mechanism by which coronaviruses reprogram host gene expression and suggest that NSP14-induced m^7^G modification is a potential therapeutic target.

## Introduction

The COVID-19 pandemic, caused by SARS-CoV-2, emerged in late 2019 and continues to pose a significant global health challenge. As a member of the *Coronaviridae* family, SARS-CoV-2 contains a positive-sense, single-stranded RNA genome of approximately 30 kb^1^. The viral genome is enclosed by a 5’ cap and 3’ poly-A tail, allowing it to function as messenger RNA (mRNA) for viral protein synthesis^2^. The 5’ end of the genome encodes two large open reading frames (ORF1a and ORF1ab), which are translated into polyproteins pp1a and pp1ab. These polyproteins are subsequently processed into 16 nonstructural proteins (NSP1-16)^3^. Each NSP plays a distinct role in viral replication and transcription, contributing to the formation of the replicase-transcriptase complex (RTC)^4–6^. Moreover, NSPs also play important roles in manipulating host gene expression and immune responses, contributing to viral replication and immune evasion^3,7^.

RNA modifications are chemical alterations to RNA molecules that play crucial roles in gene expression and RNA metabolism. To date, over 170 distinct types of RNA modifications have been identified^8^. Among these, *N*^7^-methylguanosine (m^7^G) is one of the most prevalent modifications^9^. Although m^7^G is known for its presence at the 5’ cap of mRNA, it also occurs within the internal sequences of various RNA species, including rRNA, tRNA, mRNA and non-coding RNA^10^. These modifications are catalyzed by a group of *N*7-methyltransferases (*N7*-MTases), also known as “m^7^G writers”, which methylate the guanine nucleobase at the *N*^7^ position, forming m^7^G^10^. During transcription, mRNA undergoes co-transcriptional m^7^G modification at the 5’ cap by the enzyme RNA guanine-7 methyltransferase (RNMT), which is essential for RNA stability, splicing, nuclear export and translation initiation^11,12^. In addition to capping, m^7^G modification is found in internal sequences. In tRNA, m^7^G at position G46 is installed by the METTL1-WDR4 complex and plays a critical role in RNA tertiary structure stabilization and translation fidelity^13,14^. In 18S rRNA, m^7^G at position G1639 is modified by the WBSCR22-TRM112 complex and supports ribosome biogenesis and translation initiation^15,16^. Recent studies revealed the presence of m^7^G modifications in the internal sequences of mRNA^17^. Internal m^7^G/G levels range between 0.02% and 0.05%, corresponding to approximately 800 sites across mammalian transcriptomes^17^. These internal m^7^G modifications, catalyzed by the METTL1-WDR4 complex, play important roles in translational regulation and stress responses^17–20^.

RNA modifications are prevalent strategies used by viruses to regulate both host and viral gene expression, crucial in controlling viral replication and evading host immunity^21^. Coronaviruses encode two RNA methyltransferases, NSP14 and NSP16, that modify viral RNA 5’ cap to form a functional cap structure that mimics the host mRNA cap. The 5’ cap plays a critical role in viral RNA translation and immune evasion of type I interferon (IFN-I) response^22^. NSP14 is a bifunctional enzyme with distinct *N*7-MTase and exoribonuclease (ExoN) domains^23,24^. The *N*7-MTase activity methylates the viral 5’ cap guanosine into m^7^G, forming a Cap-0 structure^25^. In contrast, NSP16 functions as a 2’-*O*-methyltransferase (2’-*O*-MTase), specifically catalyzing methylation at the 2’-*O*-ribose of the first nucleotide of the viral RNA^25^. Beyond the viral RNA capping activity, SARS-CoV-2 NSP14 plays a critical role in host RNA metabolism at the post-transcriptional level. NSP14 inhibits host translation, resulting in a reduction in IFN-I-dependent ISG induction at the protein level^26^. In addition, NSP14 inhibits mRNA export from the nucleus to the cytoplasm and alters host mRNA splicing patterns^27,28^. Notably, the *N*7-MTase activity is essential for SARS-CoV-2 replication. Recombinant SARS-CoV-2 harboring *N*7-MTase inactive mutations exhibits a significant attenuation in viral replication^29^. Moreover, inhibition of *N*7-MTase by small-molecule inhibitors restricts viral replication both *in vitro* and *in vivo*^30,31^.

In this study, we investigate the role of SARS-CoV-2 NSP14 in m^7^G modification of cellular RNA. We report that NSP14 overexpression leads to increased internal m^7^G modification in mRNA, predominantly in the nucleus. Using *N*7-MTase inactive mutants and *N*7-MTase inhibitors, we demonstrate that the *N*7-MTase activity is essential for inducing m^7^G modification. Moreover, the NSP14-NSP10 interaction enhances the NSP14-induced m^7^G modification. Notably, this m^7^G modification activity is conserved across alpha-, beta- and gamma-coronaviruses, including human coronaviruses, murine hepatitis virus (MHV) and avian infectious bronchitis virus (IBV). In addition, NSP14 induces m^7^G modification through transcriptional incorporation by hijacking cellular RNA polymerases (Pols). Specifically, NSP14 catalyzes the conversion of guanosine triphosphate (GTP) into m^7^GTP, which is then incorporated into mRNA by RNA Pol II. Moreover, NSP14-induced m^7^G modification significantly alters cellular splicing patterns. Functionally, inhibiting m^7^G modification reduces SARS-CoV-2 replication. Our results provide insights into the molecular mechanisms by which coronavirus NSP14 induces m^7^G modification in host mRNA, influencing gene expression and viral replication.

## Results

### NSP14 induces internal m^7^G modification in mRNA

Functional screenings in yeast revealed that SARS-CoV NSP14 can complement the loss of mRNA m^7^G capping activity^32^, suggesting that NSP14 modifies cellular RNA. Given the high sequence and structural homology of NSP14 proteins between SARS-CoV and SARS-CoV-2^26^, we hypothesized that SARS-CoV-2 NSP14 can also modify host RNA. To test this, the levels of RNA modification on cellular RNA were assessed by transfecting HEK293T cells with a plasmid encoding SARS-CoV-2 NSP14. Total RNA was extracted using Trizol and analyzed by liquid chromatography-tandem mass spectrometry (LC-MS/MS) to determine the levels of three RNA methylations (m^7^G, m^5^C and m^6^A). NSP14 expression was determined by immunoblot analysis (Figure 1A). The results showed endogenous RNA m^7^G, m^5^C and m^6^A modifications in the cells transfected with an empty vector control (Figure 1B). The m^6^A level was consistent with previously reported ranges^33^, whereas the levels of m^7^G and m^5^C have not been previously determined. We found that overexpression of NSP14 significantly increased m^7^G modification by 4.6-fold, whereas m^6^A and m^5^C levels remained unchanged (Figure 1B), suggesting that NSP14 induces m^7^G modification. To validate the increase in m^7^G, we performed immunoblot analysis using an m^7^G antibody, as described previously^17^. Consistent with the LC-MS/MS results, immunoblotting revealed a significant increase in m^7^G levels upon NSP14 overexpression (Figure 1C and 1D).

**Figure 1.**
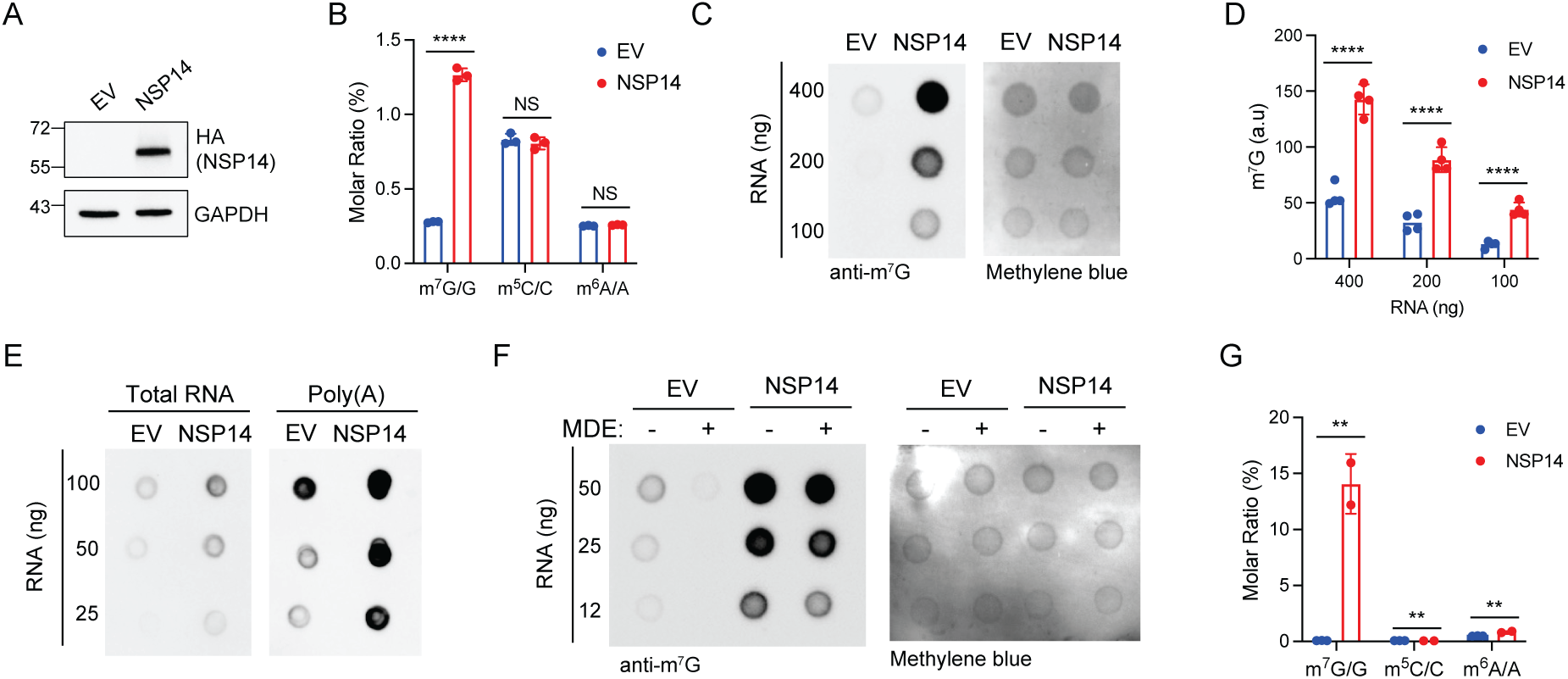
NSP14 induces internal m^7^G modification. HEK293T cells were transfected with plasmids encoding HA-tagged NSP14 or empty vector (EV) for 24 h. (A) Immunoblot analysis of HA-NSP14 expression in whole cell lysates using anti-HA and anti-GAPDH antibodies. (B) LC-MS/MS analysis of total RNA extracted from HEK293T cells transfected with EV or NSP14. RNA was Trizol-extracted, digested into nucleosides and analyzed by LC-MS/MS. The results represent the molar ratio of modified to unmodified nucleosides. (C) RNA m^7^G immunoblot analysis of total RNA from EV- or NSP14-transfected cells. Total RNA was Trizol-extracted and subjected to m^7^G immunoblotting. Methylene blue staining was used as a loading control. The intensity of m^7^G signals from four biological repeats is quantified in (D). a.u., arbitrary unit. (E) RNA m^7^G immunoblot analysis of poly(A)-selected RNA from EV- or NSP14-transfected cells. RNA was Trizol-extracted, oligo(dT)-isolated and subjected to m^7^G immunoblotting. (F) RNA m^7^G immunoblot analysis of decapped, poly(A)-selected RNA from EV- or NSP14-transfected cells. RNA was Trizol-extracted, oligo(dT)-isolated, decapped using mRNA decapping enzyme (MDE) and subjected to m^7^G immunoblotting. (G) LC-MS/MS analysis of decapped, poly(A)-selected RNA from EV- or NSP14-transfected cells. RNA was Trizol-extracted, oligo(dT)-isolated, MDE-decapped, digested into nucleosides and analyzed by LC-MS/MS. The results represent the molar ratio of modified to unmodified nucleosides. For B, D and G, data are shown as mean ± SD of two to four biological repeats. **P < 0.01, ****P < 0.001 by unpaired Student’s t test. NS, not significant.

To determine if NSP14 induces m^7^G modification in mRNA, poly(A) RNA was isolated from total RNA using oligo(dT) purification and analyzed by immunoblotting. The results showed a significant increase in m^7^G in poly(A) RNA upon NSP14 expression (Figure 1E), suggesting that NSP14 induces m^7^G in mRNA. Since mature mRNA is m^7^G-modified at the 5’ cap, we next evaluated whether NSP14-induced m^7^G can occur in internal sequences. To test this, we removed the 5’ m^7^G cap using mRNA decapping enzyme (MDE), which hydrolyzes the pyrophosphate bonds between alpha and beta phosphate moieties within the 5’-5’ triphosphate linkage, producing a 5’ monophosphate end^34^. The results demonstrated that MDE treatment significantly reduced the m^7^G levels in the control, whereas the NSP14-induced m^7^G signal remained robust (Figure 1F). These results suggest that NSP14 induces m^7^G in internal sequences. To validate the internal modification, the oligo(dT)-isolated, MDE-decapped RNA was subjected to LC-MS/MS analysis. Previous studies demonstrated an endogenous m^7^G to G ratio between 0.02% and 0.05% in cellular mRNA from various mammalian cells^17^. Consistently, in the control cells, we observed a basal level of m^7^G (Figure 1G). In contrast, NSP14 significantly increases m^7^G while both m^5^C and m^6^A levels are minimally altered, suggesting that NSP14 specifically induces internal m^7^G modification in mRNA. Together, these results demonstrate that NSP14 significantly increases m^7^G modification in the internal sequence of mRNA.

### NSP14 induces RNA m^7^G modification in the nucleus

To further study the role of SARS-CoV-2 NSP14 on RNA methylation, we performed immunofluorescence analysis in HeLa cells transfected with either GFP or GFP-tagged NSP14. Confocal microscopy analysis demonstrated a significant increase in m^7^G signal intensity in cells expressing GFP-NSP14 compared to cells expressing GFP (Figure 2A). This increase in m^7^G signal was consistently observed under both low and high contrast settings. Quantitative analysis of m^7^G fluorescence intensity across individual GFP-positive cells confirmed a significant elevation of m^7^G levels in both HeLa and HEK293T cells expressing GFP-NSP14, relative to GFP controls (Figure 2B).

**Figure 2.**
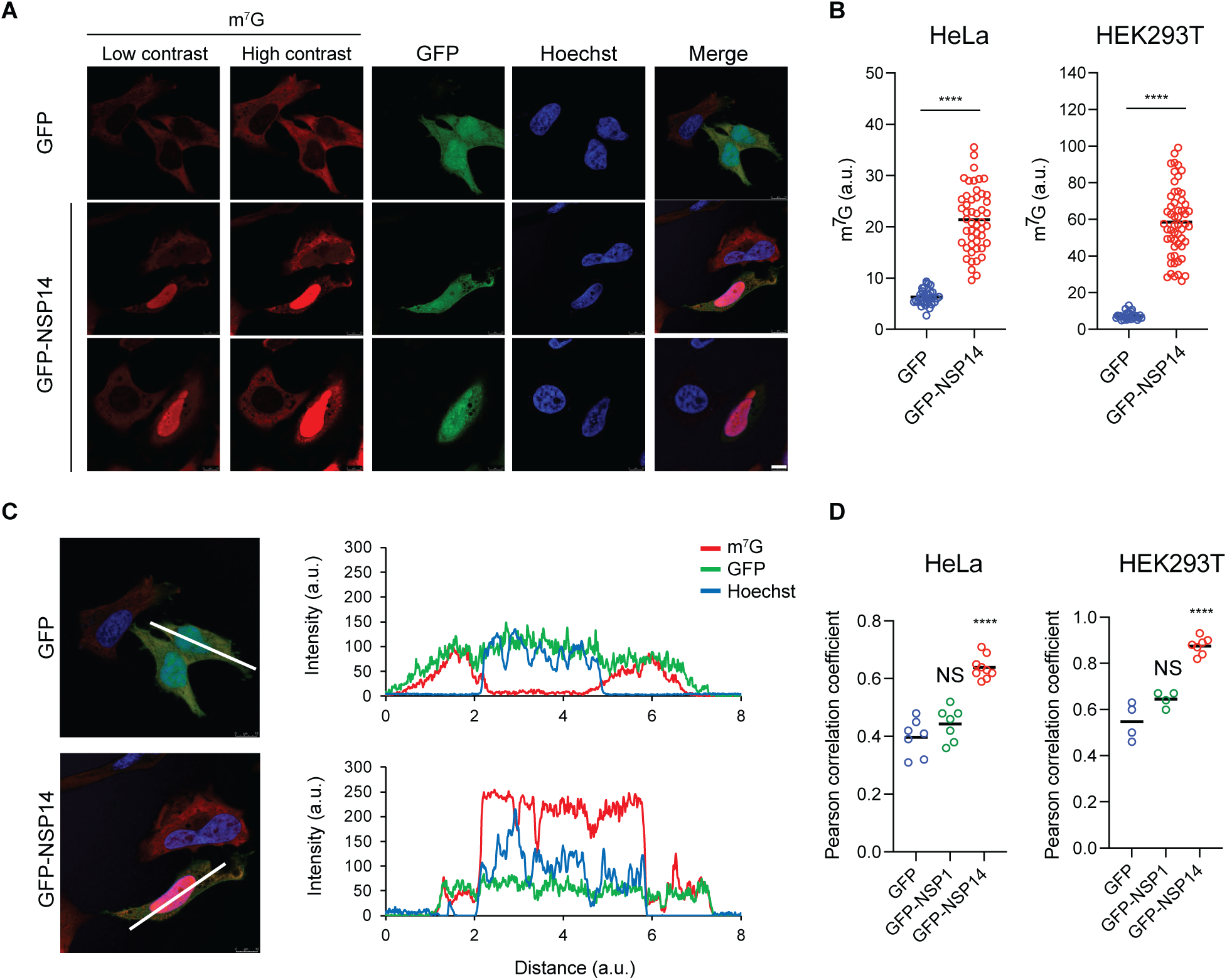
NSP14 induces m^7^G modification in the nucleus. (A) Confocal microscopy analysis of HeLa cells transfected with plasmids encoding GFP or GFP-tagged NSP14 (GFP-NSP14). After 24 h of transfection, cells were fixed and stained with m^7^G antibody (red) and Hoechst (blue). Scale bar, 10 µm. (B) Quantification of m^7^G fluorescence intensity from individual GFP-positive cells shown in (A). Data represent the mean fluorescence intensities of GFP and GFP-NSP14-transfected HeLa cells (left) and HEK293T cells (right). ****P < 0.001 by unpaired Student’s t test. a.u., arbitrary unit. (C) Fluorescence intensity profiles of m^7^G, GFP and Hoechst signals along the white marked lines in the merged images shown in (A). (D) Quantification of colocalization between m^7^G and the nucleus (Hoechst). Data represent Pearson correlation coefficients for the colocalization analysis of GFP, GFP-NSP1 and GFP-NSP14-transfected HeLa cells (left) and HEK293T cells (right). Data are shown as the mean of four to eight repeats. ****P < 0.001 by unpaired Student’s t test. NS, not significant.

Notably, the NSP14-induced m^7^G signals colocalize with the nuclear DNA marker Hoechst (Figure 2A). To determine the subcellular localization of the m^7^G signal, we analyzed fluorescence intensity profiles along lines drawn through representative cells. In GFP-expressing cells, the m^7^G signal was low and diffusely distributed in the cytoplasm, whereas in GFP-NSP14-expressing cells, the m^7^G signal was sharply elevated and colocalized with the nuclear marker Hoechst (Figure 2C). The intensity plots showed a clear overlap between m^7^G and Hoechst fluorescence in NSP14-expressing cells, but not in GFP controls, indicating that NSP14-induced m^7^G modification predominantly occurs in the nucleus. To quantify this spatial association, we performed colocalization analysis between m^7^G and Hoechst signals using Pearson correlation coefficients. In both HeLa and HEK293T cells, GFP-NSP14 expression significantly increased the nuclear colocalization of m^7^G compared to GFP or GFP-NSP1 controls (Figure 2D). Notably, GFP-NSP1, another viral nonstructural protein, did not show significant changes in m^7^G levels or nuclear localization, suggesting a specific effect of NSP14 on nuclear RNA methylation.

Despite the partial cytoplasmic localization of NSP14, the m^7^G signal was enriched in the nucleus, suggesting that NSP14 may act indirectly through host nuclear factors or facilitate modification of nuclear RNA substrates. Together, these results demonstrate that NSP14 increases m^7^G RNA methylation, with a predominant nuclear localization pattern, supporting its potential role in manipulating host RNA processing machinery during viral infection.

### *N*7-MTase activity is required to induce m^7^G modification

NSP14 is a versatile enzyme with two distinct catalytic domains. The N-terminal 3’ to 5’ ExoN domain removes incorrectly incorporated nucleotides, which plays a critical role in maintaining high RNA replication fidelity^35^. The C-terminal domain exhibits *N*7-MTase activity, which is responsible for modifying viral RNA at the 5’ cap guanosine, catalyzing the formation of the m^7^G cap structure (cap-0)^29^. Notably, mutations that inactivate either domain significantly impair viral replication^29,35^. To determine the role of these activities in RNA m^7^G modification, we used two NSP14 mutants, H268A (M2) and D331A/G333A (M4), which disrupt the ExoN and *N*7-MTase activities, respectively^26^. We transfected HEK293T cells with either WT NSP14 or the mutants. Consistent with our previous study^26^, both mutants showed higher protein expression levels than WT (Figure 3A). To assess their roles in RNA modification, total RNA was extracted from the transfected cells and analyzed by m^7^G immunoblotting. The results showed that both mutants lost the m^7^G modification activity (Figure 3B). Immunofluorescence analysis further demonstrated that both mutant proteins failed to induce m^7^G modification in the nucleus (Figure 3C and 3D). To further study the role of *N*7-MTase in internal m^7^G modification of mRNA, RNA was extracted from HEK293T transfected with NSP14 and M4 mutant, oligo(dT)-isolated, MDE-decapped and subjected to LC-MS/MS analysis. The results demonstrated that the *N*7-MTase mutant failed to induce internal m^7^G modification in mRNA (Figure 3E).

**Figure 3.**
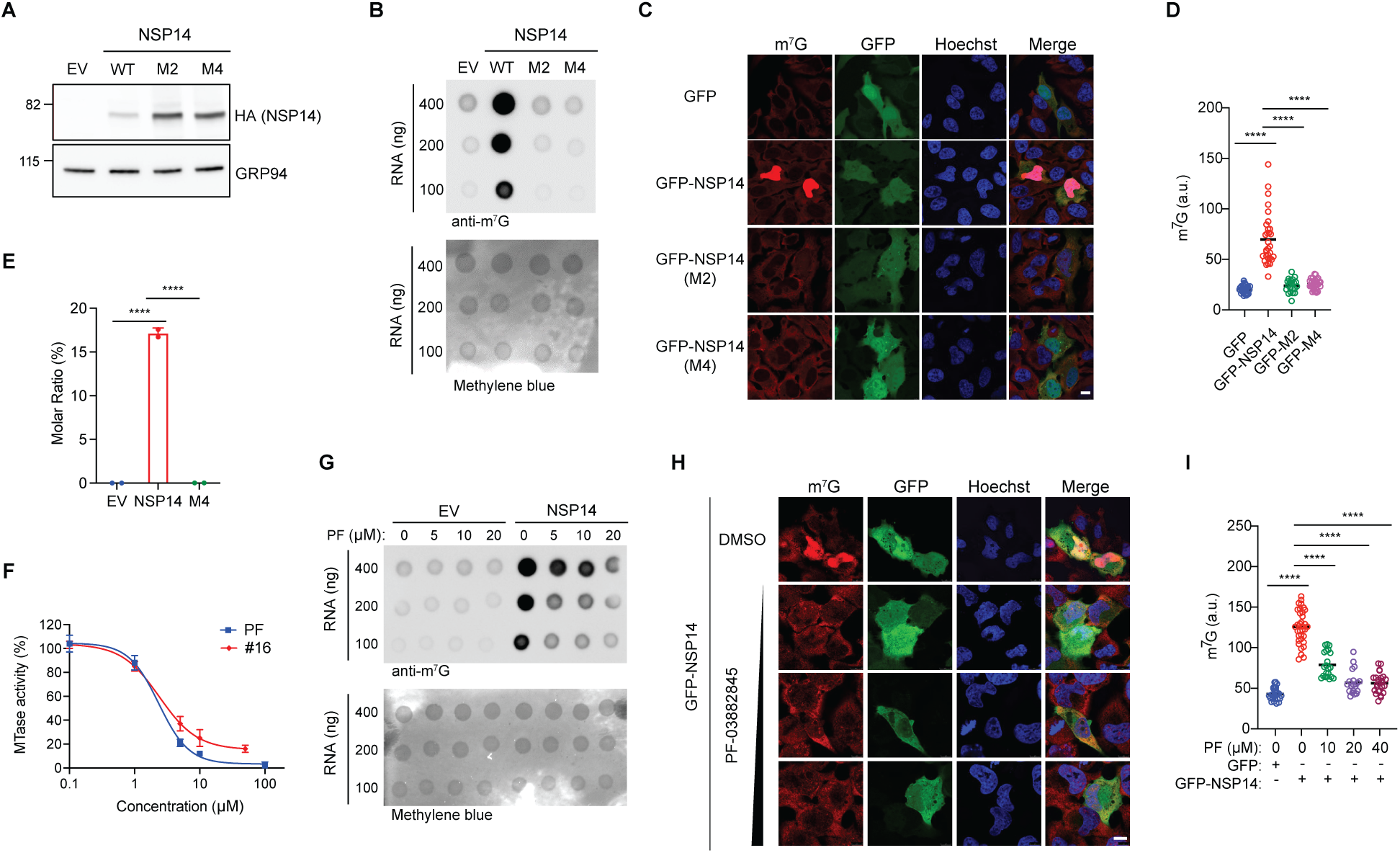
*N*7-MTase activity is required to induce m^7^G modification. (A) Immunoblot analysis of HEK293T cells transfected with EV, wildtype (WT) NSP14, the exonuclease mutant (M2) or the *N*7-methyltransferase mutant (M4). After 24 h of transfection, cell lysates were subjected to immunoblot analysis with anti-HA and anti-GRP94 antibodies. (B) RNA m^7^G immunoblot analysis of total RNA from transfected cells shown in (A). After 24 h of transfection, total RNA was Trizol-extracted and subjected to m^7^G immunoblotting. Methylene blue staining was used as a loading control. (C) Confocal microscopy analysis of HeLa cells transfected with plasmids encoding GFP or GFP-tagged WT and NSP14 mutants. After 24 h of transfection, cells were fixed and stained with anti-m^7^G antibody (red) and Hoechst (blue). Scale bar, 10 µm. (D) Quantification of m^7^G fluorescence intensity from individual GFP-positive cells shown in (C). Data are shown as mean fluorescence intensity. ****P < 0.001 by unpaired Student’s t test. a.u., arbitrary unit. (E) LC-MS/MS analysis of m^7^G modification in decapped, poly(A)-selected RNA from HEK293T cells transfected with EV, NSP14 or M4 for 24 h. RNA was Trizol-extracted, oligo(dT)-isolated, MDE-decapped, digested into nucleosides and analyzed by LC-MS/MS. The results represent the molar ratio of modified to unmodified nucleosides. Data are shown as mean ± SD of two biological repeats. ****P < 0.001 by unpaired Student’s t test. (F) *In vitro* MTase activity assay using the recombinant SARS-CoV-2 NSP14-NSP10 complex and GTP as a substrate. The *N*7-MTase inhibitors, PF-03882845 (PF) and Compound 16 (#16), were used to inhibit *N*7-MTase activity. Data are shown as mean ± SD of three biological repeats. (G) Dose-dependent inhibition of NSP14-induced m^7^G RNA modification in HEK293T cells treated with increasing concentrations of PF. Cells were transfected with EV or WT NSP14 and treated with PF for 24 h. Total RNA was Trizol-extracted and subjected to m^7^G immunoblotting. (H) Confocal microscopy images of HeLa cells transfected with GFP-NSP14 and treated with PF (10, 20, 40 µM). After 24 h, cells were fixed and stained with m^7^G antibody (red) and Hoechst (blue). Scale bar, 10 µm. (I) Quantification of m^7^G fluorescence intensity from individual GFP-positive cells shown in (H). Data are shown as mean fluorescence intensity. ****P < 0.001 by unpaired Student’s t test.

To confirm the role of *N*7-MTase, we utilized two *N*7-MTase inhibitors of SARS-CoV-2 NSP14, PF-03882845 and Compound 16 (#16)^30,36^. Using *in vitro* MTase assays, we demonstrated that PF-03882845 and #16 reduce the MTase activity of NSP14 with IC^50^ values of 2.35 μM and 2.39 μM, respectively (Figure 3F). To further investigate the role of *N*7-MTase activity in m^7^G modification, we treated HEK293T and HeLa cells with PF-03882845 and transfected the cells with NSP14. Using m^7^G immunoblotting, we showed that PF-03882845 treatment significantly reduced RNA m^7^G modification in NSP14-transfected cells in a dose-dependent manner (Figure 3G). Similar results were observed in HeLa cells transfected with GFP-NSP14 using immunofluorescence staining (Figure 3H and 3I). Consistent with the literature^25^, we confirmed that PF-03882845 exhibited no detectable cytotoxicity under the experimental conditions (Figure S1). Together, these findings suggest that *N*7-MTase activity is required for internal m^7^G modification.

### NSP10-NSP14 interaction increases internal m^7^G modification

SARS-CoV NSP14 protein forms a protein complex with the viral protein NSP10, which enhances its enzymatic activities^24,37^. Previous studies revealed the interaction in the SARS-CoV-2 homologs and that the formation of the complex significantly enhances the activities of NSP14 in exoribonuclease and translational regulation^26,38^. To study the role of NSP10 in NSP14-induced m^7^G modification, we co-transfected HEK293T cells with plasmids encoding NSP10 and NSP14. Protein levels of NSP10 and NSP14 were assessed by immunoblotting, which confirmed that co-expression with NSP10 significantly enhanced NSP14 expression (Figure 4A), consistent with our previous findings^2^. Using m^7^G immunoblotting, we found that co-transfection of NSP10 and NSP14 increased m^7^G modification (Figure 4B). To determine the role of NSP10-NSP14 interaction in m^7^G modification, we used NSP10 mutants that have compromised NSP10-NSP14 interaction. We previously showed that two NSP10 mutations at the protein-protein interface led to reduced (K43A) or no interaction (H80A) with NSP14^2^. Consistently, compromising NSP10-NSP14 interaction resulted in reduced NSP14 expression (Figure 4C). Notably, the impaired interaction led to reduced m^7^G modification (Figure 4D). Together, these results suggest that the formation of the NSP10-NSP14 complex stabilizes NSP14 and increases its protein level and the RNA m^7^G modification.

**Figure 4.**
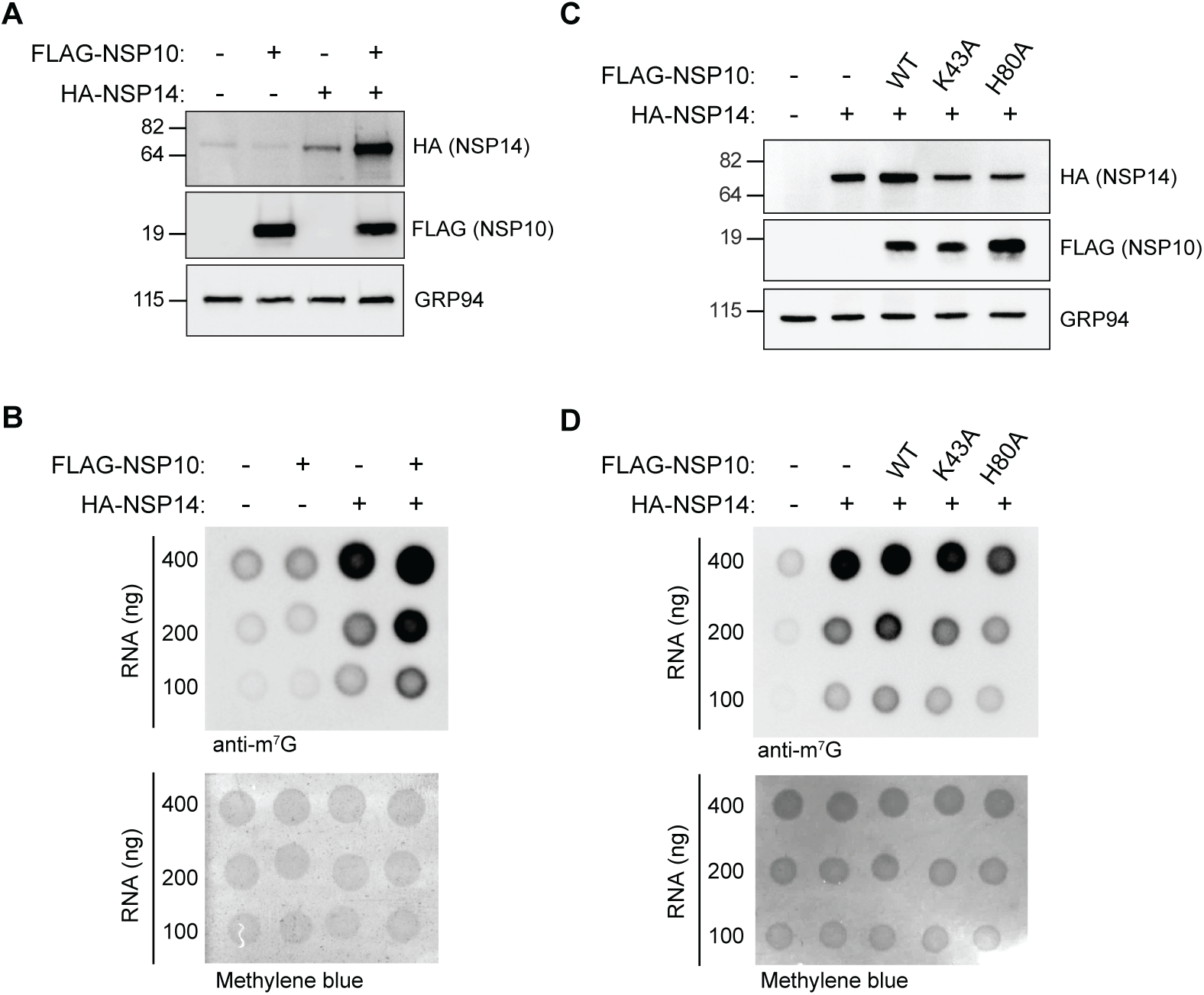
The NSP10-NSP14 interaction enhances internal m^7^G modification. (A) Immunoblot analysis of HEK293T cells co-transfected with plasmids encoding HA-tagged NSP14 (HA-NSP14) and FLAG-tagged NSP10 (FLAG-NSP10) for 24 h. Cell lysates were subjected to immunoblot analysis with anti-HA, anti-FLAG and anti-GRP94 antibodies. (B) RNA m^7^G immunoblot analysis of HEK293T cells transfected with the indicated plasmids. After 24 h, total RNA was Trizol-extracted and subjected to m^7^G immunoblotting and methylene blue staining. (C) Immunoblot analysis of HEK293T cells transfected with plasmids encoding HA-NSP14, FLAG-NSP10 or its mutants (K43A and H80A) for 24 h. Cell lysates were subjected to immunoblot analysis with anti-HA, anti-FLAG and anti-GRP94 antibodies. (D) RNA m^7^G immunoblot analysis of HEK293T cells transfected with the indicated plasmids. After 24 h, total RNA was Trizol-extracted and subjected to m^7^G immunoblotting and methylene blue staining.

### The internal m^7^G modification activity is conserved in coronaviruses

The *N*7-MTase activity of NSP14 is essential for the viral RNA 5’ end modification in human coronaviruses (HCoV), which plays a critical role in viral replication and evasion of host immune responses^29,39^. Disruption of *N*7-MTase activity, through either mutations or chemical inhibitors, significantly impairs viral replication^29–31^. Sequence analysis of CoV NSP14 proteins reveals high sequence homology (Figure S2). Notably, the SAM-binding motif (DxGxPxG/A) in the *N*7-MTase domain^29^ is conserved within CoV (Figure 5A). To study whether this *N*7-MTase activity extends to RNA m^7^G modification, we transfected HEK293T cells with NSP14 from different coronaviruses. First, we examined highly pathogenic human beta-coronaviruses, including SARS-CoV, SARS-CoV-2 and MERS-CoV. The protein expression levels were determined by immunoblotting (Figure 5B). Using m^7^G immunoblotting, we found significant increases in RNA m^7^G modification induced by these NSP14 homologs (Figure 5C), suggesting a conserved RNA modification activity.

**Figure 5.**
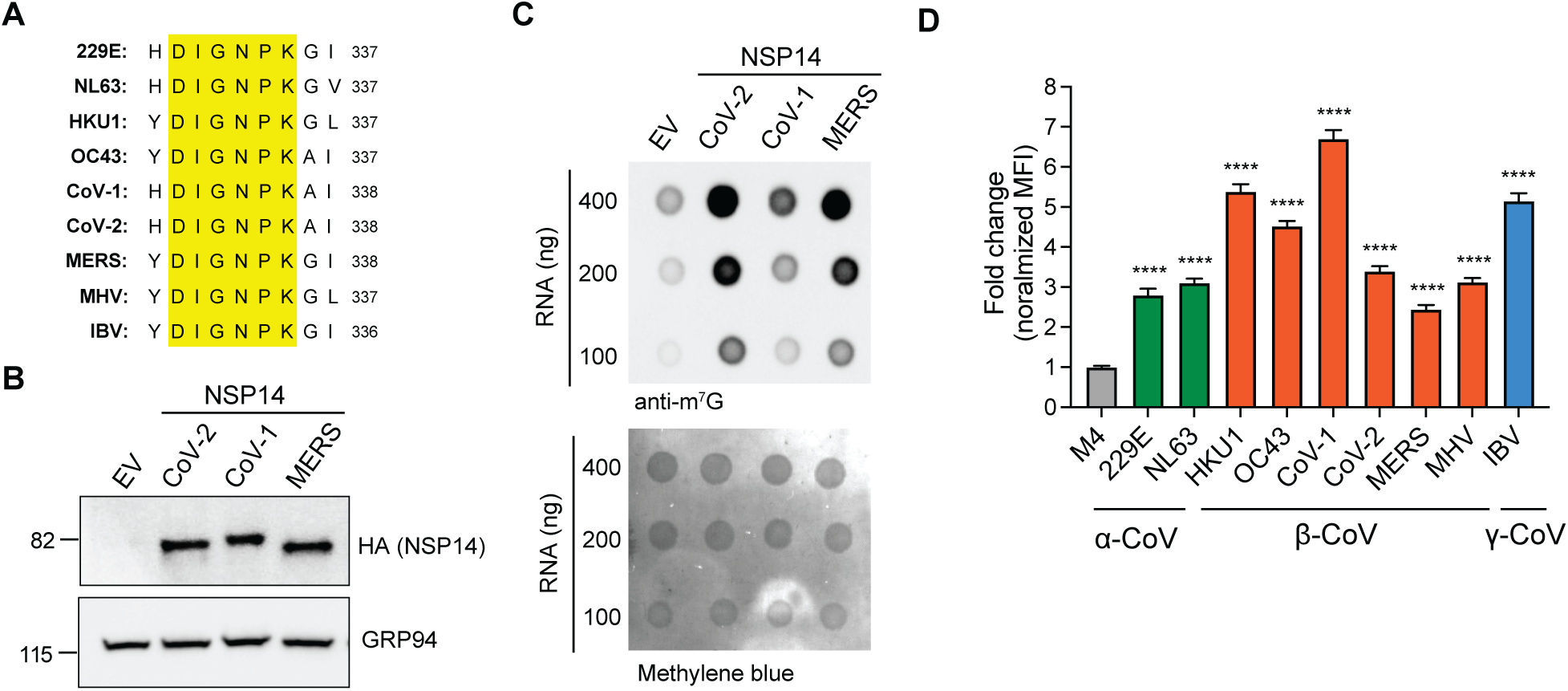
The internal m^7^G activity is conserved in coronaviruses. (A) Protein sequence alignment of coronavirus NSP14 at the *N*7-MTase active site. Conserved amino acid residues are highlighted in yellow. Human alphacoronavirus (HCoV-229E [229E], HCoV-NL63 [NL63]), human betacoronaviruses (HCoV-HKU1 [HKU1], HCoV-OC43 [OC43], MERS-CoV [MERS], SARS-CoV [CoV-1], SARS-CoV-2 [CoV-2]), murine betacoronavirus (mouse hepatitis virus [MHV]) and avian gammacoronavirus (infectious bronchitis virus [IBV]). (B) Immunoblot analysis of HEK293T cells transfected with plasmids encoding different HA-tagged coronavirus NSP14 proteins for 24 h. Cell lysates were subjected to immunoblot analysis with anti-HA and anti-GRP94 antibodies. (C) RNA m^7^G immunoblot analysis of HEK293T cells transfected with the indicated plasmids. After 24 h, total RNA was Trizol-extracted and subjected to m^7^G immunoblotting and methylene blue staining. (D) Flow cytometry analysis of HEK293T cells transfected with plasmids encoding HA-tagged NSP14 proteins from different coronaviruses for 24 h. Cells were fixed, stained with anti-m^7^G and anti-HA antibodies and analyzed by flow cytometry. Data are presented as mean ± SD of three to nine biological repeats. ****P < 0.001 by unpaired Student’s t test.

To further screen the RNA modification activity, we developed a flow cytometry approach using m^7^G antibody (Figure S3A). In brief, HEK293T cells were transfected with plasmids encoding HA-tagged NSP14. RNA m^7^G modification at the single-cell level was assessed by intracellular staining with m^7^G antibody. Flow cytometry analysis revealed that, at the single-cell level, SARS-CoV-2 NSP14 significantly increased the m^7^G signal compared to the *N*7-MTase mutant (Figure S3B). We applied this method to assess the m^7^G modification activity of NSP14 from coronaviruses. The results confirmed that NSP14 proteins from highly pathogenic human betacoronaviruses exhibited m^7^G modification activity (Figure 5D). Moreover, we found similar activity in NSP14 from the common cold human alpha-and betacoronaviruses associated with mild to moderate, self-limiting respiratory diseases, including HCoV-229E, HCoV-NL63, HCoV-HKU1 and HCoV-OC43 (Figure 5D). Interestingly, this activity is also conserved in a murine betacoronavirus, mouse hepatitis virus (MHV) and an avian gammacoronavirus, avian infectious bronchitis (IBV) (Figure 5D). Together, these results suggest that the m^7^G modification is a conserved activity of NSP14 proteins across coronaviruses.

### NSP14 catalyzes the conversion of GTP into m^7^GTP

Internal m^7^G modifications are typically catalyzed by a group of *N*7-MTases, commonly known as m^7^G writers, which directly methylate RNA at specific sites^10^. However, despite its catalytic potential, NSP14 alone does not appear sufficient to directly induce internal m^7^G modification. Localization studies demonstrated that NSP14 is present in both the cytoplasm and nucleus^40–42^, but NSP14-induced m^7^G modification is predominantly localized in the nucleus (Figure 2). This result indicates that while NSP14 is essential for m^7^G modification, it cannot independently perform this function, implying the involvement of additional nuclear factor(s) that facilitate the modification process. Moreover, previous studies showed that SARS-CoV NSP14 can catalyze the conversion of GTP to m^7^GTP *in vitro*^43^. In addition, cellular expression of SARS-CoV-2 NSP14 is associated with increased intracellular m^7^GTP^27^. Therefore, we hypothesized that RNA Pols catalyze NSP14-induced m^7^G modification by incorporating m^7^GTP into RNA.

To study the structural basis of this activity, we first employed homology modeling and molecular docking to study the binding interactions of SARS-CoV-2 NSP14 with nucleotide triphosphates (NTPs), including GTP, ATP, CTP and UTP. We used two structural templates, SARS-CoV-2 NSP14 bound to the cap analog m^7^GpppG (PDB 7QIF) and the SARS-CoV-1 NSP14-NSP10 complex with GpppA (PDB 5C8S). To model SARS-CoV-2-specific interactions in the SARS-CoV structure, we substituted homologous amino acids. Per-residue interaction energies were calculated to assess the contribution of individual residues, and docking simulations were repeated five times. The molecular docking results showed that, in both structures of NSP14 and the NSP14-NSP10 complex, the complementary function (CF) score of GTP had a larger negative value than other NTPs, suggesting that GTP has the strongest binding affinity among the tested NTPs, followed by ATP, UTP and CTP (Figure 6A). We also found that this specificity was largely driven by a π–π stacking interaction between the guanine base of GTP and the aromatic side chain of F426 (Figure 6B). In contrast, pyrimidine-based NTPs like UTP and CTP lacked this interaction and fell outside the binding cavity (Figure S4A). Moreover, the *N*7-methylated guanine of m^7^GTP was positioned outside the binding cavity (Figure S4B), and docking analyses indicate that m^7^GTP bound less favorably than GTP (Figure 6A), consistent with its function as a product released from the NSP14 active site.

**Figure 6.**
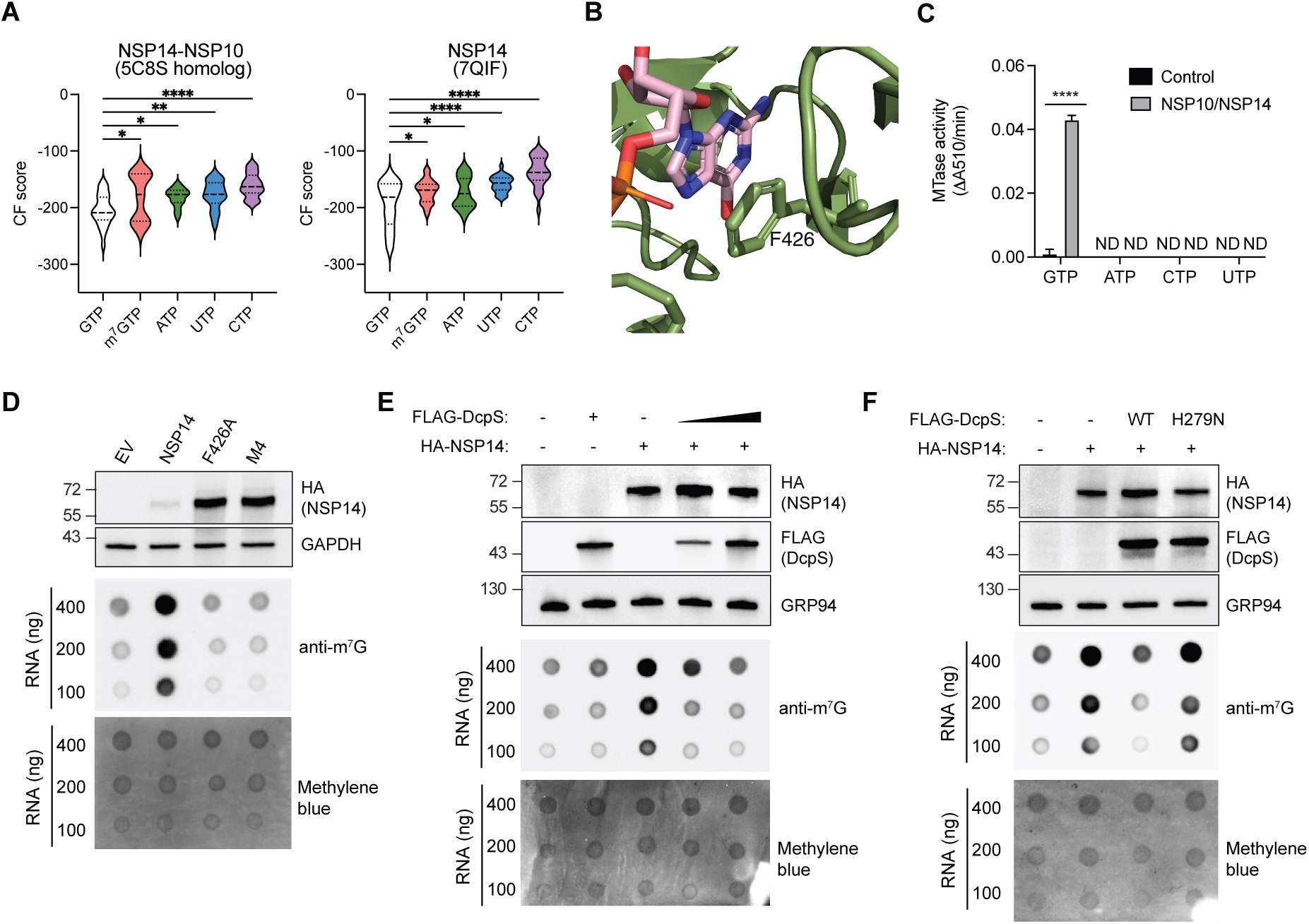
NSP14 methylates GTP to generate m^7^GTP for RNA modification. (A) Molecular docking analysis of SARS-CoV-2 NSP14 with various nucleotide triphosphates, illustrating differential binding affinities. The complementary function (CF) score is a docking metric that reflects the predicted binding affinity; more negative values indicate stronger binding. Data are shown as violin plots representing the top five poses from five independent docking runs (25 poses in total). *P < 0.05, **P < 0.01, ****P < 0.001 by unpaired Student’s t test. (B) Structural model highlighting the π-π stacking interaction between the guanine base of GTP (pink) and residue F426 of NSP14 (green). (C) *In vitro* methyltransferase (MTase) activity assay of the NSP14-NSP10 complex using nucleotide triphosphates. Data are presented as mean ± SD of three biological repeats. ****P < 0.001 by unpaired Student’s t test. ND, not detected. (D) Protein and RNA m^7^G immunoblot analyses of HEK293T cells transfected with WT, F426A and M4 mutant NSP14 for 24 h. Cell lysates were subjected to immunoblot analysis with anti-HA and anti-GAPDH antibodies. Total RNA was Trizol-extracted and subjected to m^7^G immunoblotting and methylene blue staining. (E) Protein and RNA m^7^G immunoblot analyses of HEK293T cells co-transfected with plasmids encoding HA-tagged NSP14 (HA-NSP14) and FLAG-tagged DcpS (FLAG-DcpS) for 24 h. (F) Protein and RNA m^7^G immunoblot analyses of HEK293T cells co-transfected with plasmids encoding HA-NSP14 and WT or inactive DcpS mutant (H279N) for 24 h.

To validate the modeling results, we examined the catalytic activity of NSP14 in synthesizing m^7^GTP. We conducted an *in vitro* methylation assay using NTPs. The results demonstrated that NSP14 methylated GTP but not ATP, UTP and CTP (Figure 6C). The identity of the methylated product was confirmed as m^7^GTP by mass spectrometric analysis (Figure S5). To test the role of F426 in RNA m^7^G modification, we transfected HEK293T cells with plasmids encoding WT and F426A NSP14 and assessed m^7^G using immunoblotting. The results demonstrated that F426A significantly reduced NSP14-induced m^7^G (Figure 6D), suggesting a critical role for F426 in substrate recognition and catalysis. Notably, F426 is conserved among coronaviruses, supporting its importance for viral replication.

To further confirm the involvement of m^7^GTP in m^7^G modification in cells, we employed the scavenger mRNA decapping enzyme DcpS, which can degrade cellular m^7^GTP by converting it into m^7^GMP^27^. We co-transfected HEK293T cells with plasmids encoding NSP14 and DcpS and assessed m^7^G using immunoblotting. The results demonstrated that DcpS significantly reduced NSP14-induced m^7^G in a dose-dependent manner (Figure 6E). Moreover, we also generated an inactive mutant of DcpS, H279N^44^. The results showed that H279N failed to inhibit NSP14-induced m^7^G (Figure 6F), suggesting that m^7^GTP is required for the NSP14-induced m^7^G. Together, these molecular modeling and experimental results demonstrate that NSP14 catalyzes the synthesis of m^7^GTP from GTP. This product is then required, likely through incorporation by RNA Pols, for internal RNA m^7^G modification.

### NSP14 induces internal m^7^G via RNA polymerase-dependent incorporation

To study the role of RNA Pol activity in NSP14-induced m^7^G, we used transcription inhibitors to block RNA synthesis in cells. Since these inhibitors can suppress transcription from plasmid DNA, we instead transfected HEK293T cells with *in vitro* transcribed mRNA encoding either WT NSP14 or the *N*7-MTase inactive mutant (M4). Consistent with results from plasmid transfection, NSP14 mRNA transfection induced m^7^G modification, whereas the M4 mutant failed to exhibit this activity (Figure 7A). To inhibit transcription, HEK293T cells were treated with actinomycin D, a broad-spectrum transcription inhibitor, followed by transfection with NSP14 mRNA. The results showed that actinomycin D significantly reduced m^7^G modification (Figure 7B), indicating an important role for active transcription in NSP14-induced m^7^G. Given that NSP14 increases m^7^G levels in mRNA (Figure 1E and 1G), we next examined the effects of selectively inhibiting RNA Pol II activity using α-amanitin. The results demonstrated that α-amanitin treatment significantly inhibited m^7^G modification (Figure 7C). In contrast, treatment with CX-5461, an RNA Pol I inhibitor, had no effect on m^7^G modification (Figure 7D). Additionally, we confirmed that the transcription inhibitors exhibited no detectable cytotoxicity under the experimental conditions (Figure S6A and S6B). The effectiveness of the inhibitors was confirmed by RT-qPCR analysis of genes transcribed by each RNA Pol: 47S rRNA (Pol I), Myc (Pol II) and U6 snRNA (Pol III) (Figure S6C and S6D).

**Figure 7.**
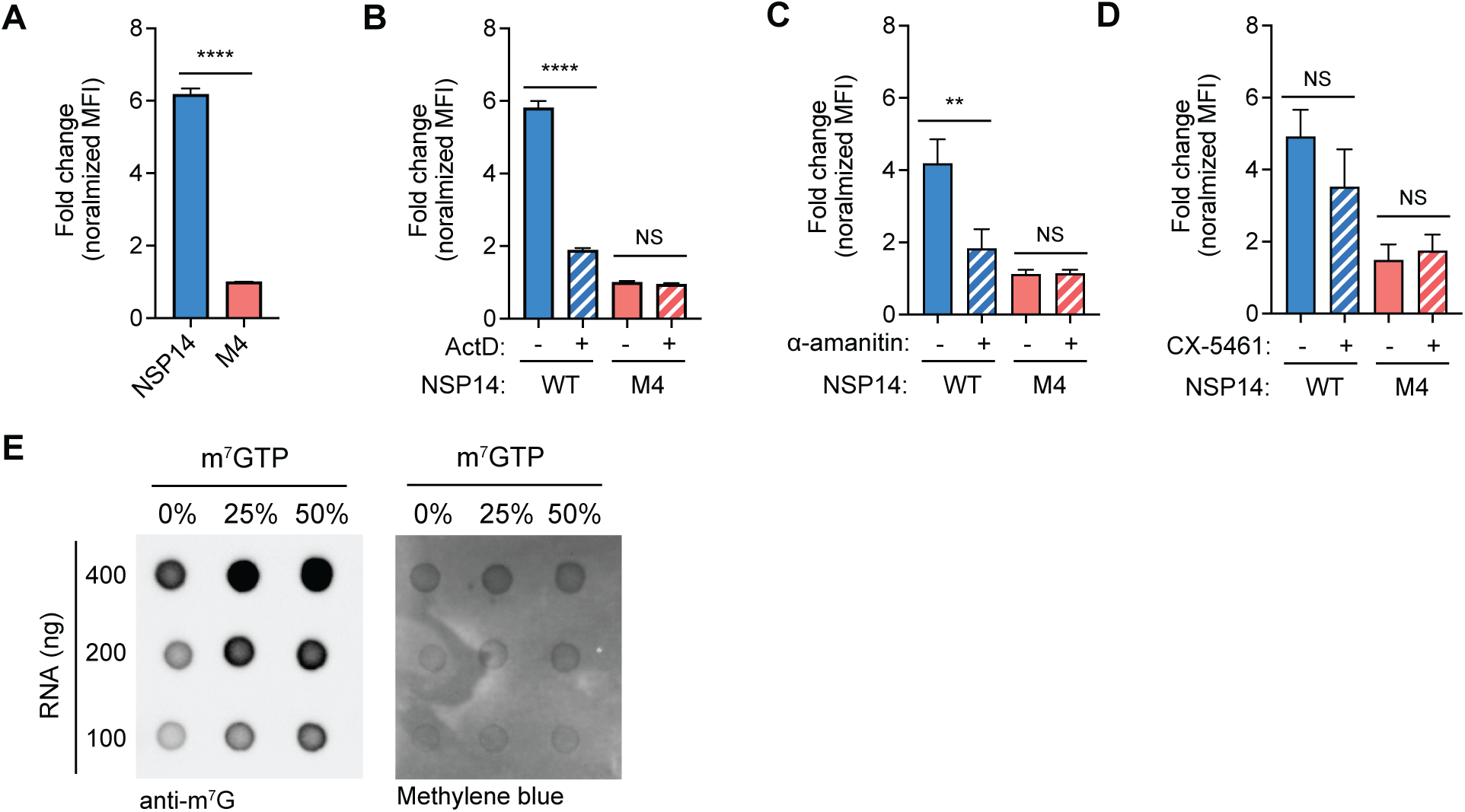
RNA polymerase II incorporates m^7^GTP into RNA. (A) Flow cytometry analysis of HEK293T cells transfected with *in vitro*-transcribed RNA encoding NSP14 or M4 for 24 h. Cells were fixed, stained with anti-m^7^G and anti-HA antibodies and analyzed by flow cytometry. Data are presented as mean ± SD of three biological repeats. ****P < 0.001 by unpaired Student’s t test. (B-D) Flow cytometry analysis of HEK293T cells transfected with *in vitro* transcribed RNA encoding NSP14 or M4 and treated with (B) actinomycin D (ActD), (C) α-amanitin or (D) CX-5461 for 24 h. Cells were fixed, stained with anti-m^7^G and anti-HA antibodies and analyzed by flow cytometry. Data are presented as mean ± SD of three biological repeats. **P < 0.01, ****P < 0.001 by unpaired Student’s t test. NS, not significant. (E) RNA m^7^G immunoblot analysis of *in vitro*-transcribed RNA. RNA was synthesized using HeLa nuclear extract from a CMV promoter-driven DNA fragment, with m^7^GTP supplemented at indicated ratios. The resulting RNA was analyzed by m^7^G immunoblotting and methylene blue staining. The background m^7^G signals observed at 0% m7GTP are due to endogenous RNA in HeLa nuclear extract.

To study the role of RNA Pol II in m^7^GTP incorporation into mRNA, we used a human *in vitro* transcription system with HeLa nuclear extract. In brief, we employed a DNA template containing a cytomegalovirus (CMV) promoter, which drives expression of the enhanced green fluorescent protein (EGFP) gene in a Pol II-dependent manner. To assess the incorporation of m^7^GTP, we supplemented the *in vitro* transcription reaction with m^7^GTP and subsequently analyzed the *in vitro* transcribed RNA using m^7^G immunoblotting. The results showed that m^7^GTP was incorporated into RNA by RNA Pol II (Figure 7E). Together, these results indicate that NSP14 induces internal m^7^G modification through a mechanism that depends on RNA Pol II-mediated incorporation of NSP14-generated m^7^GTP into mRNA.

### RNA m^7^G modification induces alternative splicing

Betacoronaviruses, including SARS-CoV-2, induce alternative splicing in host RNA^45^. Previous studies identified that SARS-CoV-2 NSP14 significantly induces alternative splicing, primarily promoting intron retention^27,28^. Notably, the *N*7-MTase activity of NSP14 is required for this effect^27,28^. To study the role of m^7^G modification in alternative splicing, we first examined how NSP14 affects the formation of partial novel and novel splice junctions^46^. Partial novel splice junctions are splicing events in which one splice site (either the 5’ donor or 3’ acceptor) matches a known exon boundary, but the other site is unannotated. These events likely reflect mispairing between canonical and cryptic splice sites. Novel splice junctions, in which both splice sites are unannotated, represent entirely aberrant splicing events and indicate profound disruption of normal splicing regulation. RNA-Seq analysis showed that expression of NSP14 induced a significant increase in partial novel and novel splice junctions, without affecting canonical splicing events (Figure 8A). In contrast, cells expressing an *N*7-MTase-deficient mutant of NSP14 (M4) showed no such changes, suggesting that the *N*7-MTase activity of NSP14 is essential for altering host splicing patterns.

**Figure 8.**
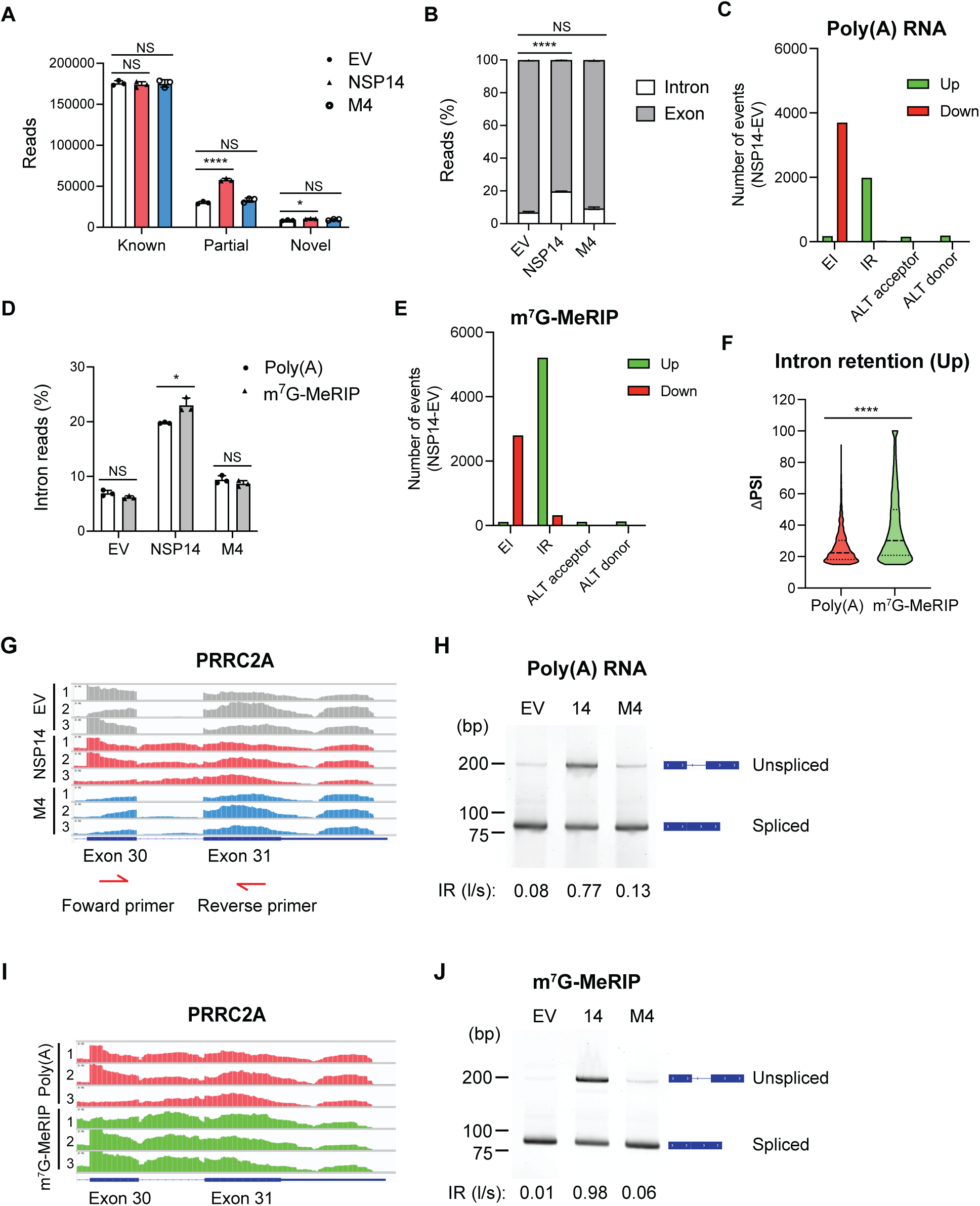
m^7^G modification induces alternative splicing. (A) Splice junction analysis of poly(A) RNA from HEK293T cells transfected with empty vector (EV), wildtype NSP14 (WT) or the *N*7-MTase mutant (M4) for 24 h. Poly(A) RNA was isolated using oligo(dT) from total RNA and subjected to RNA-Seq analysis. NSP14 expression increased the number of partial novel and novel splicing junctions but not known junctions. (B) RSeQC read distribution analysis of poly(A) RNA from HEK293T cells transfected with EV, NSP14 or M4 for 24 h. Poly(A) RNA was isolated as described in (A). (C) Quantification of alternative splicing events in poly(A) RNA between NSP14- and EV-transfected cells. EI, exon inclusion. IR, intron retention. ALT, alternative. (D) RSeQC read distribution analysis of m^7^G-enriched mRNA from HEK293T cells expressing EV, NSP14 or M4 for 24 h. Poly(A) RNA was isolated using oligo(dT) from total RNA, then MDE-decapped, immunoprecipitated by anti-m^7^G antibody to isolate m^7^G-MeRIP RNA. Both poly(A) RNA and m^7^G-MeRIP RNA were subjected to RNA-Seq. (E) Quantification of alternative splicing events in m^7^G-enriched mRNA between NSP14- and EV-transfected cells. (F) Analysis of the level of NSP14-induced intron retention. Violin plots show the distribution of ΔPSI (percent spliced-in index) values for the up-regulated intron retention events in poly(A) RNA versus m^7^G-MeRIP RNA. ΔPSI values reflect the change in splicing inclusion levels between NSP14- and EV-transfected cells. NSP14 expression led to significantly higher ΔPSI values in m^7^G-modified RNA compared to poly(A) RNA. (G) RNA-Seq tracks of PRRC2A showing increased intron reads between exons 30 and 31 in cells expressing NSP14. Poly(A) RNA from HEK293T cells expressing EV, NSP14 or M4 were isolated as described in (A). Forward and reverse primers used for RT-PCR to detect intron retention are indicated. (H) RT-PCR analysis of intron retention in poly(A) RNA from HEK293T cells transfected with EV, NSP14 or M4 for 24 h. Poly(A) RNA was reverse-transcribed and amplified by PCR. The resulting PCR products were analyzed by DNA gel electrophoresis. The intron retention ratio of unspliced “long” fragments to spliced “short” fragments (IR(l/s)) was quantified using ImageJ. (I) RNA-Seq tracks of PRRC2A comparing poly(A) RNA and m^7^G-MeRIP RNA, showing higher intron retention in m^7^G-enriched RNA. RNA was extracted and subjected to m^7^G-MeRIP-Seq as shown in (D). (J) RT-PCR analysis of intron retention in m^7^G-MeRIP RNA from HEK293T cells transfected with EV, NSP14 or M4, as described in (H). m^7^G-MeRIP RNA, rather than poly(A) RNA, was used for RT-PCR. For A, B, D and F, data are presented as mean ± SD of three biological repeats. *P < 0.05, ****P < 0.001 by unpaired Student’s t test. NS, not significant.

Among the major types of alternative splicing, previous studies demonstrated that NSP14 increases intron retention and reduces exon inclusion^27,28^, both of which contribute to an increased level of introns. Consistently, we showed that NSP14, but not the M4 mutant, significantly increased intron reads in poly(A) RNA (Figure 8B), accompanied by elevated intron retention and reduced exon inclusion (Figure 8C). To further study the role of m^7^G in alternative splicing, we performed m^7^G-methylated RNA immunoprecipitation (m^7^G-MeRIP) to enrich internal m^7^G-modified mRNA. In brief, mRNA was oligo(dT)-isolated, MDE-decapped and subjected to RNA immunoprecipitation using m^7^G antibody. RNA-Seq analysis revealed that NSP14 induced an increase in intron reads in internal m^7^G-modified RNA (Figure 8D). Notably, the intron levels were significantly higher in internal m^7^G-modified RNA compared to poly(A) RNA, suggesting an important role for m^7^G in modulating splicing patterns. Moreover, NSP14 induced more intron retention events in internal m^7^G-modified RNA compared to poly(A) RNA (Figure 8C vs 8E). In addition, NSP14-induced m^7^G modifications not only increased intron retention events but also elevated the levels of retention (Figure 8F).

Intron retention can generate aberrant protein isoforms and abolish protein function. To study the biological significance of NSP14-induced intron retention, we performed gene ontology (GO) analysis on genes with increased intron retention (Figure S7). The results revealed that affected genes are strongly enriched for processes related to genome stability and RNA metabolism, including pathways involved in DNA replication and repair, cell cycle regulation and RNA processing. At the cellular level, these genes localize predominantly to nuclear compartments associated with chromatin organization, splicing and protein quality control. Functionally, they are enriched for enzymatic and regulatory activities acting on DNA and RNA, including helicases, transcription factors and chromatin-modifying enzymes. Together, these results indicate that NSP14-induced intron retention broadly disrupts nuclear programs that coordinate genome maintenance with RNA processing, suggesting a mechanism by which the virus reshapes host gene expression to favor infection.

To validate the changes in alternative splicing, we performed reverse transcription followed by PCR to quantify intron retention levels in selected target genes. We focused on three genes, PRRC2A, ATG101 and RPS27A, which exhibited increased intron retention upon NSP14 expression, whereas SSBP3 served as a negative control (Figure S8A). For each gene, we designed a set of primers targeting a specific intron that displayed increased retention (Figure 8G and S8B). These primers span adjacent exon boundaries, producing ∼200 bp amplicons from unspliced transcripts, whereas shorter products are amplified from fully spliced mRNA (Figure 8H and S9). Using poly(A) RNA as templates, we found that NSP14 increased the abundance of long fragments compared to EV and M4, indicating an increase in unspliced RNA levels (Figure 8H and Figure S9). Given that the m^7^G-MeRIP-Seq results revealed a higher level of intron retention (Figure 8E and 8F), we further analyzed the unspliced RNA levels using m^7^G-MeRIP RNA (Figure 8I and Figure S10). For example, m^7^G-enriched RNA showed an increase in the intron reads between Exon 30 and 31 of PRRC2A compared to poly(A) RNA (Figure 8I). Consistently, RT-PCR results showed an increase in unspliced RNA levels, as indicated by an increase in the IR(l/s) ratio from 0.77 to 0.98 (Figure 8H vs 8J). Together, these findings demonstrate that NSP14 induces intron retention in m^7^G-modified RNA.

To further validate the role of NSP14-induced m^7^G modification in alternative splicing, we utilized DcpS to reduce internal m^7^G modification levels. Cells were co-transfected with the plasmids encoding NSP14 and DcpS. The alternative splicing patterns of target genes were analyzed using RT-PCR. Co-transfection with DcpS significantly reduced NSP14-induced intron retention, restoring splicing patterns to those observed in control conditions (Figure S11). Together, these findings demonstrate that NSP14 induces alternative splicing in m^7^G-modified RNA.

### Inhibiting NSP14-induced alternative splicing reduces SARS-CoV-2 replication

A previous study identified a direct correlation between SARS-CoV-2 viral load and splicing isoform distribution in patient samples^45^. Higher viral loads are associated with significant shifts in alternative splicing patterns in SARS-CoV-2 infection, suggesting that inducing alternative splicing enhances viral replication. To study the role of NSP14-induced alternative splicing in SARS-CoV-2 replication, Vero E6 cells were infected with SARS-CoV-2 at a multiplicity of infection (MOI) of 1 for 24 hours with *N*7-MTase or RNA Pol inhibitors. The cell culture medium was harvested to determine viral titers using focus-forming assays (FFA). The results demonstrated that the *N*7-MTase inhibitor PF-03882845 significantly reduced SARS-CoV-2 replication (Figure 9A), consistent with previous studies^30^. This result suggests the important role of the *N*7-MTase activity of NSP14 in viral replication.

**Figure 9.**
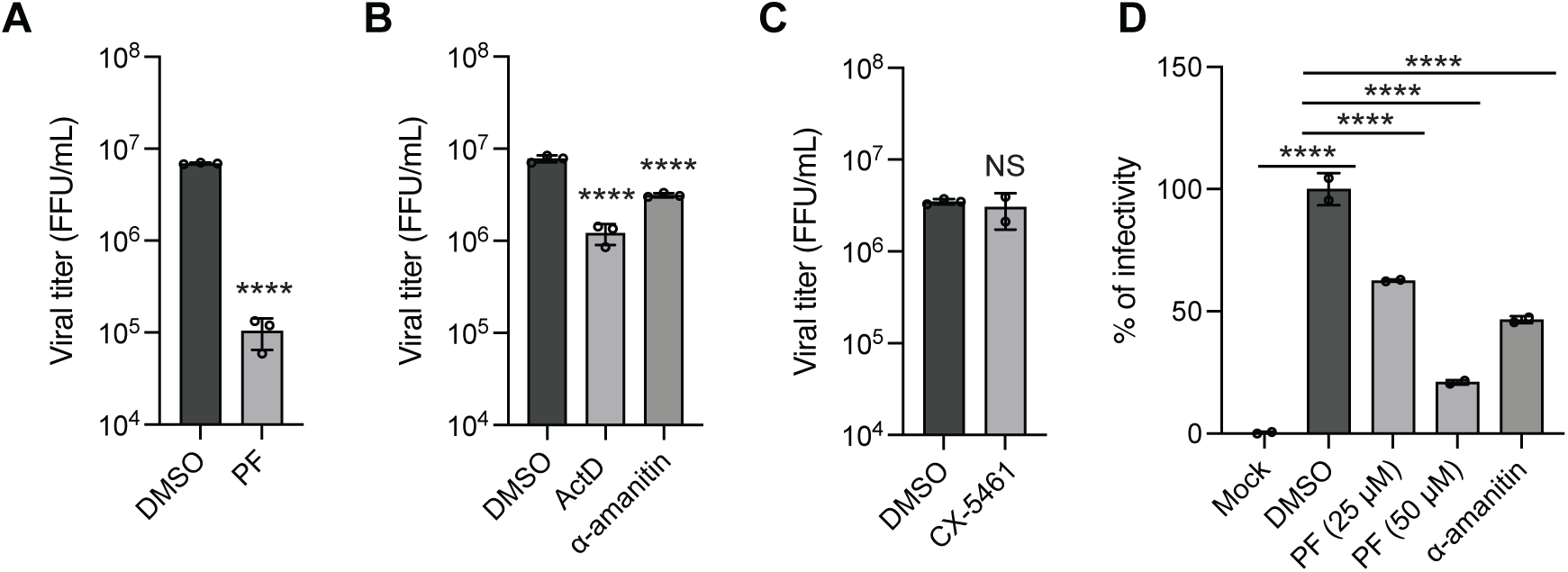
*N*7-MTase and transcription inhibitors reduce SARS-CoV-2 replication. (A-C) Vero E6 cells were infected with SARS-CoV-2 (WA1/2020) at a multiplicity of infection (MOI) of 1 and treated with the indicated inhibitors for 24 h. Cells were treated with (A) the NSP14 *N*7-MTase inhibitor PF-03882845 (PF) and (B-C) transcription inhibitors (actinomycin D [ActD], α-amanitin, CX-5461). Supernatants were collected, and viral titers were determined using focus-forming assays. FFU, focus-forming unit. (D) A549-Ace2-Dpp4-Tmprss2 cells were infected with SARS-CoV-2 XEC variant at an MOI of 1 and treated with the indicated inhibitors for 24 h. Supernatants were collected, and viral titers were determined immunofluorescence-based viral replication assay. Data are presented as mean ± SD of two to three biological repeats. ****P < 0.001 by unpaired Student’s t test. NS, not significant.

As inhibition of the *N*7-MTase activity of NSP14 affects both viral RNA 5’ capping and host RNA m^7^G modification, we next focused on the role of NSP14-induced host m^7^G modification in viral replication. Since internal m^7^G modification is incorporated by RNA Pols (Figure 7), we employed transcription inhibitors to selectively disrupt host m^7^G modification. The results showed that actinomycin D treatment significantly inhibited viral replication (Figure 9B). Notably, inhibition of RNA Pol II with α-amanitin also led to a significant reduction in viral replication (Figure 9B). In contrast, inhibition of RNA Pol I with CX-5461 did not affect viral replication (Figure 9C). We confirmed that PF-03882845 and transcription inhibitors exhibited no detectable cytotoxicity under the experimental conditions (Figure S12).

To further assess the dependence of SARS-CoV-2 variants on RNA Pol II activity, we examined a clinical isolate of the XEC variant, an Omicron subvariant that accounted for an estimated 45% of U.S. cases in early December 2024. Consistently, inhibition of RNA Pol II with α-amanitin significantly inhibits XEC replication (Figure 9D). Together, these results demonstrate that NSP14-induced alternative splicing and internal m^7^G modification are critical for efficient SARS-CoV-2 replication, and that disruption of these processes can substantially impair viral replication.

## Discussion

Coronavirus NSP14 plays canonical roles in viral RNA capping and proofreading via its *N*7-MTase and ExoN domains, respectively. Both enzymatic activities are indispensable for efficient viral replication, suggesting the essential role of NSP14 in the coronavirus life cycle^29,35^. Beyond its functions on viral RNA, recent studies have revealed a versatile role for NSP14 in regulating host RNA metabolism. SARS-CoV-2 NSP14 alters host RNA splicing, inhibits RNA nuclear export and suppresses host protein synthesis^26–28^. Here, we uncovered a novel function of NSP14 in reshaping host RNA epitranscriptomics and splicing patterns through internal m^7^G modification. The m^7^G modification activity is specific to the internal sequences of polyadenylated mRNA and is enriched in the nuclear compartment. Notably, this activity is conserved across multiple coronaviruses, including SARS-CoV-1, SARS-CoV-2, MERS-CoV and other human and animal coronaviruses. Using an *N*7-MTase mutant and NSP14 inhibitors, we demonstrated that *N*7-MTase activity is essential for m^7^G modification. Mechanistically, NSP14 catalyzes the conversion of GTP into m^7^GTP, which is then incorporated into host mRNA by RNA Pol II. This reveals a novel mechanism by which viruses exploit host transcriptional machinery to induce non-canonical RNA modifications. Importantly, this study also links NSP14-induced m^7^G modification to alternative splicing regulation. Inhibiting either the *N*7-MTase activity or m^7^G incorporation significantly impaired SARS-CoV-2 replication, suggesting that SARS-CoV-2 hijacks the host epitranscriptomic landscape to reprogram gene expression to favor viral replication. Together, this study provides new insight into the importance of RNA modifications in virus-host interactions and identifies NSP14 as a key driver of coronavirus-mediated epitranscriptomic changes.

SARS-CoV-2 NSP14 modulates host RNA metabolism by suppressing translation, blocking nuclear export and remodeling the transcriptome^26–28^. These transcriptomic changes affect thousands of genes and include altered mRNA abundance and localization, disrupted splicing patterns and upregulation of circular RNAs, many of which are implicated in innate immunity. Notably, the *N*7-MTase activity of NSP14 is essential for these changes in RNA metabolism^26–28^. This activity, typically responsible for viral RNA capping, appears to have an unanticipated regulatory role in host gene expression. Moreover, through its *N*7-MTase activity, NSP14 activates inosine 5’-monophosphate dehydrogenase 2 (IMPDH2), the rate-limiting enzyme in the de novo guanine nucleotide biosynthesis, leading to elevated GTP levels that drive metabolic and transcriptional changes, including NF-κB pathway activation and *CXCL8* induction^28^. Additionally, NSP14 disrupts mRNA splicing and nuclear export by targeting the host nuclear cap-binding complex (CBC), interfering with 5’ cap recognition^27^. This impairs the processing of pre-mRNAs and blocks their export to the cytoplasm. In this study, we demonstrated that NSP14 induces m^7^G modification in mRNA, which installs a methyl group at the *N*7 position of the guanine in the internal sequences. This modification can potentially alter RNA structure and RNA-protein interactions, contributing to changes in splicing, nuclear export and translational regulation. Together, these findings reveal the *N*7-MTase domain of NSP14 as a multifunctional effector that orchestrates transcriptional reprogramming, impairs mRNA maturation and supports viral immune evasion by disrupting host RNA metabolism at multiple levels.

Internal RNA modifications play critical roles in cellular function and human diseases. These modifications have been implicated in tumorigenesis, immune evasion and reduced drug sensitivity, highlighting their potential as biomarkers and therapeutic targets in cancer and other diseases^47–50^. In viral infections, m^6^A modifications regulate viral replication and host immune responses^51^. Flavivirus infection induces m^6^A modifications in a subset of host mRNAs, regulating their expression through alternative splicing and translational regulation, resulting in a cellular environment that supports viral replication^52^. SARS-CoV-2 infection induces dynamic changes in the m^6^A methylome, with increased m^6^A methylation detected in both viral and host mRNA^53,54^. For m^7^G modifications, stress conditions have been shown to induce internal mRNA modification that influences RNA translation and stability^55,56^. Furthermore, a higher expression of most m^7^G-related genes in COVID-19 compared to non-COVID-19 patients has been found, suggesting a role in viral pathogenesis^57^. Here, we reported a novel function of SARS-CoV-2 NSP14 in internal m^7^G modification and alternative splicing. Importantly, inhibition of NSP14’s *N*7-MTase activity or blocking m^7^G incorporation impairs SARS-CoV-2 replication, reinforcing the idea that the virus manipulates host epitranscriptome to establish a cellular environment conducive to viral replication. By reshaping the host mRNA landscape at the level of RNA modifications and splicing, NSP14 emerges not only as a key factor in viral replication but also as a modulator of host gene expression. Together, our findings expand the known repertoire of coronavirus-host interactions and identify the NSP14-induced internal m^7^G pathway as a critical axis of epitranscriptomic reprogramming with implications for viral pathogenesis, immune evasion and potential therapeutic targeting.

RNA modifications are typically installed posttranscriptionally by specific enzymes known as “writers”^58^. These proteins recognize consensus RNA sequences or structural motifs and catalyze the corresponding chemical reactions^58^. Several cellular m^7^G writers are known to catalyze internal m^7^G modifications^59^. 18S rRNA is m^7^G modified at G1639 by the WBSCR22-TRM112 complex, whereas a subset of tRNAs is m^7^G modified at G46 by the METTL1-WDR4 complex^13–16^. These modifications play critical roles in RNA structure and translation initiation and fidelity^13–16^. mRNA also undergoes internal m^7^G modification by the METTL1-WDR complex, which plays important roles in translational regulation and stress responses^17–20^. Coronavirus NSP14 represents the first known viral protein that can induce internal m^7^G modification. Rather than acting as a conventional m^7^G writer, NSP14 methylates GTP into m^7^GTP, which is subsequently incorporated into mRNA by RNA Pol II. Additionally, through activation of IMPDH2, NSP14 enhances *de novo* guanosine nucleotide synthesis, elevating GTP levels that fuel m^7^GTP production^28^. This novel RNA modification mechanism reveals an unprecedented virus-host interaction strategy used by coronaviruses to reprogram host RNA metabolism. Interestingly, transcription-coupled incorporation of nucleotide analogs is widely exploited in both basic research and therapeutic applications. For example, 4-thiouridine (4-SU), 5-ethynyluridine (5-EU) and 5-bromouridine (5-BrU) are incorporated into nascent RNA transcripts and used to study

RNA synthesis and turnover^60^. In cancer therapy, drugs like 5-fluorouracil (5-FU) are incorporated into cellular RNA, interfering with RNA processing and function^61^. In antiviral therapies, nucleoside analogs such as remdesivir, favipiravir, ribavirin and molnupiravir are incorporated into viral RNA by viral RNA-dependent RNA Pols. This incorporation leads to premature chain termination, lethal mutagenesis or disruption of RNA synthesis, which ultimately inhibits viral replication^62–65^. Notably, ddhCTP, a naturally occurring antiviral nucleoside analog produced by the interferon-stimulated protein viperin, is also incorporated into viral RNA, where it halts viral RNA chain elongation^66^. These examples highlight how both viruses and host cells exploit RNA Pol-mediated incorporation of modified nucleotides as a strategy to modulate RNA function.

RNA modifications are key regulators of RNA structure and RNA-protein interactions^67^. Among these, m^6^A is well known for its role in weakening A-U base pairing through methylation at the N6 position of adenosine, which alters base-pairing dynamics at the Watson-Crick interface^68^. This structural destabilization exposes single-stranded regions that facilitate binding by proteins such as hnRNP-C and hnRNP-G^69–71^, an effect termed the “m^6^A switch.” m^6^A also directly recruits m^6^A reader proteins like the YTH family, influencing splicing, transcription, mRNA export, translation and decay^67^. Consistently, depletion of METTL3, the key m^6^A methyltransferase, leads to splicing defects in hundreds of genes. Other modifications, such as *N*^1^-methyladenosine (m^1^A) and *N*^1^-methylguanosine (m^1^G), also impact RNA folding by disrupting non-canonical Hoogsteen base pairs, leading to A-form RNA destabilization^72^. Here, we found that NSP14 induces a dramatic increase in the internal mRNA m^7^G/G ratio, from 0.05% to 14%. m^7^G involves methylation at the *N*7 position of guanine nucleobase, introducing a positive charge without affecting Watson–Crick base pairing directly. However, this modification alters the electron distribution of guanosine, potentially disrupting non-canonical interactions. RNA G-quadruplexes (rG4s), structures stabilized by Hoogsteen hydrogen bonding between guanines, are particularly sensitive to m^7^G. rG4s play regulatory roles in pre-mRNA splicing, where they can enhance or suppress the use of nearby splice sites by recruiting specific RNA-binding proteins or remodeling local RNA structure^73^. Mimicking internal m^7^G modification with 7-deaza-deoxyguanosine (DAG) disrupts key hydrogen bonds, leading to rG4 destabilization^74^. By modulating rG4 stability, NSP14-induced internal m^7^G modifications may act as structural switches, analogous to the m^6^A switch, leading to alternative splicing.

An intriguing aspect of virus-host interactions is the potential evolutionary impact that viruses may have exerted on the architecture of host immune genes, particularly type I interferons (IFN-I). One of the defining features of mammalian and avian IFN-I genes is their intronless structure^75^, with the notable exception of IFN-κ, which retains a single intron^76^. In contrast, ancestral IFN-I genes in fish and amphibians exhibit a multi-exon structure composed of five exons and four introns^77,78^. It is hypothesized that the retroposition of intron-containing IFN-I transcripts gave rise to the intronless IFN-I genes found in amniotes, likely occurring during the evolutionary transition of vertebrates from aquatic to terrestrial environments^78^. This phylogenetic divergence raises an important question in evolutionary immunology: did selective pressure from pathogens favor the emergence of intronless IFN-I genes in amniotes^79^? Our findings that SARS-CoV-2 NSP14 promotes intron retention in host transcripts via internal m^7^G modification suggest that intron-containing mRNAs may be especially vulnerable to virus-induced post-transcriptional disruption. Molecular clock analyses suggest that the most recent common ancestor of coronaviruses could date back as far as 190-489 million years^80^, potentially overlapping with early vertebrate or amniote evolution. Importantly, many other ancient virus families, such as rhabdoviruses and paramyxoviruses, have co-evolved with vertebrates for hundreds of millions of years and are known to interfere with host RNA processing and splicing. Given that IFN-I serves as a central antiviral effector, its intronless architecture may have evolved to ensure rapid, splicing-independent expression in response to viral immune invasion, escaping viral manipulation of RNA processing. Together, these observations support a model in which viruses that disrupt host RNA splicing may have contributed to the evolution of IFN-I genes, favoring intronless gene structures that are resistant to splicing-based antagonism.

## Limitations

Our study reveals a novel mechanism by which coronavirus NSP14 induces internal m^7^G modification and alters host mRNA splicing. Several limitations should be acknowledged. First, while we demonstrate that m^7^GTP incorporation is RNA Pol II–dependent, the specific sequence or structural features that guide internal m^7^G placement remain unclear. Future studies using crosslinking or high-resolution mapping will be required to define the m^7^G consensus sites and their transcriptomic distribution. Second, although we show that altered mRNA splicing correlates with increased intron retention and impaired antiviral gene expression, the functional consequences of specific splice isoform changes, such as their stability, translation or contribution to viral replication and immune evasion, were not evaluated. Third, although inhibition of NSP14-induced m^7^G modification and alternative splicing reduces viral replication, the precise downstream host pathways or genes driving this antiviral effect are still unclear. Functional studies linking specific splicing changes to viral fitness or immune evasion will be crucial to understanding the broader implications of NSP14 activity. Lastly, the potential interplay between NSP14-induced m^7^G modification and other epitranscriptomic marks (e.g., m^6^A, m^5^C) remains an open question that requires further investigation.

## METHOD DETAILS

### Plasmids

Viral protein constructs were generated as described previously^26^. DcpS was cloned from HEK293T cDNA into the pCAGGS-FLAG vector. NSP10, NSP14 and DcpS mutants were generated using site-directed mutagenesis. All primers used for molecular cloning and qRT-PCR were synthesized by Integrated DNA Technologies (IDT). Plasmids were purified using Zymo Research kits.

### Cells culture

HEK293T, HeLa and Vero E6 cells were cultured in Dulbecco’s modified Eagle’s media (DMEM; Corning) supplemented with 10% fetal bovine serum (FBS; Corning). For DNA plasmid and mRNA transfection, cells were plated 24 h before transfection and transfected using Lipofectamine 2000 according to the manufacturer’s instructions. For inhibitor experiments, cells were pre-treated with the indicated concentrations of inhibitors for 24 h prior to transfection or infection. Inhibitors remained in the culture medium throughout the post-transfection or post-infection period unless otherwise specified.

### Viruses and cell culture infections

The SARS-CoV-2 virus (isolate USA-WA1/2020) was obtained from the BEI Resources repository. All infection experiments were conducted under biosafety level 3 (BSL-3) containment as described previously^81,82^. Vero E6 cells were mock-infected or infected with the virus at a multiplicity of infection (MOI) of 1 in serum-free DMEM for 1 h. After removing inoculum, plates were overlaid with 1.2% methylcellulose containing 2% FBS for. 24 h. Cells were fixed with 4% paraformaldehyde or 10% formaldehyde for 1 h at room temperature, PBS washed three times and stained with anti-nucleocapsid antibody for SARS-CoV-2 (1:1000) for 1 h at 37 °C. Cells were then washed three times with PBS and stained with HPR-conjugated secondary antibody for 1 h at 37 °C. Foci were visualized by peroxidase substrate, after washing the cells twice with PBS. Viral titers were quantified as fluorescent focus units (FFU) per mL.

The SARS-CoV-2 isolate XEC (V7447) was isolated from a nasopharyngeal clinical sample at the Hackensack Meridian Health Center for Discovery and Innovation (HMH-CDI). Viruses were propagated in Vero E6-Tmprss2 cells in a growth medium consisting of DMEM and 2% heat-inactivated FBS. Virus infections were performed in A549-Ace2-Dpp4-Tmprss2 (ADT) cells for 24 h at 37 °C in infection medium (DMEM, 10% FBS)^83^. Immunofluorescence-based viral replication assay and cytotoxic assay were performed as described previously^83^. In brief, immunofluorescence staining for viral nucleocapsid protein (NP) was performed to determine viral replication activity. Infected cells were 4% PFA-fixed, PBS-washed, Triton X-100 (0.1%)/PBS permeabilized and blocked with blocking solution (0.1% Triton X-100 and 0.2% BSA in PBS). Cells were stained with NP-specific antibodies, Alexa-594-conjugated secondary antibodies and DAPI. Images were acquired with Cytation C10 and processed using the Gen 5 Image Software (Agilent BioTek). The toxicity of the compounds was measured using CellTiter Glo with a Tecan Infinite M200PRO plate reader to measure the cell viability by luminescence.

### Protein immunoblotting

Cells were lysed in Laemmli buffer and boiled for 10 min. Cell lysates were separated by sodium dodecyl sulfate polyacrylamide gel electrophoresis (SDS-PAGE) with 4-15% Mini-PROTEAN TGX gels (Bio-Rad) and transferred onto PVDF membranes (Millipore). Membranes were blocked in 5% BSA/Tris-buffered saline with 0.1% Tween 20 (TBST) for 1 h at room temperature, incubated with primary antibodies in 2% BSA/TBST overnight at 4°C, followed by incubation with HRP-conjugated antibodies in 2% BSA/TBST for 1 h at room temperature. Signal detection was performed using the SuperSignal West Pico PLUS kit (Thermo Scientific) in a ChemiDoc MP Imaging System (Bio-Rad).

### RNA extraction, poly(A) RNA purification and decapping reaction

Cells were harvested and washed with DPBS buffer. Total RNA was extracted using TRIzol reagent (Thermo Scientific) and subsequently purified with Direct-zol RNA MiniPrep (Zymo Research). mRNA was enriched by two rounds of poly(A) purification using oligo(dT)25 magnetic beads (New England Biolabs). RNA concentrations were measured using Nanodrop spectrophotometer. For mRNA decapping reaction, poly(A) RNA was treated with mRNA Decapping Enzyme (New England Biolabs) according to the manufacturer’s protocol and purified using Oligo Clean & Concentrator (Zymo Research).

### RNA Immunoblotting (m^7^G dot blot)

RNA immunoblotting was performed as described previously with minor modifications^17^. In brief, RNA was serially diluted and spotted on a nylon membrane, followed by UV-crosslinking. Membranes were blocked with 5% BSA/TBST for 1 h at room temperature and incubated with anti-m^7^G antibody (RN017M, MBL) in 2% BSA/TBST. After three washes with 1 x TBST, membranes were incubated with HRP-conjugated secondary antibodies in 0.05% non-fat milk/TBST for 1 hour at room temperature. Signal detection was performed using the SuperSignal West Pico PLUS kit (Thermo Scientific) in a ChemiDoc MP Imaging System (Bio-Rad).

### Confocal immunofluorescence microscopy

HeLa cells were seeded onto glass coverslips and transfected. After 24 h, the cells were washed with PBS and fixed with 4% paraformaldehyde/PBS for 10 min at room temperature, followed by incubation in cold methanol for 10 minutes on ice. Cells were then incubated with 70% ethanol for 10 min, followed by 1M Tris-HCl (pH 8) for 5 min. Permeabilization and blocking were performed with 1% BSA in 2x SSC buffer containing 0.1% Triton X-100 for 30 min. anti-m^7^G antibody (1:200) in blocking buffer was applied overnight at 4°C. After washing three times with 2x SSC buffer, cells were incubated with anti-mouse IgG Fab2 (Thermo Fisher, A21237, 1:100) for 1 h at room temperature in blocking buffer. Following three washes with 2x SSC buffer, cells were stained with Hoechst 1:1,000 in 2x SSC buffer for 10 min and washed twice with 2x SSC buffer. Coverslips were mounted using ProLong Gold Anti-Fade Reagent (Thermo Fisher), and imaging was conducted using a Leica SP8 confocal microscope. Images were analyzed using ImageJ.

### RT-PCR and qRT-PCR

RNA was extracted from cells using TRIzol and reverse-transcribed into cDNA. For qRT-PCR, RNA was quantified using Luna Universal One-Step RT-qPCR Kit (New England Biolabs). Analyses were performed using an Mx3000P real-time PCR system (Stratagene). For RT-PCR, splicing patterns were analyzed using primers targeting intron-exon boundaries by Luna Universal One-Step RT-qPCR Kit (New England Biolabs). DNA products were analyzed by 12% TBE polyacrylamide gel electrophoresis (PAGE). For experiments involving transcriptional inhibitors, total RNA was extracted using TRIzol reagent and reverse-transcribed into cDNA. Quantification of RNA levels was performed using Luna® Universal qPCR Master Mix (M3003, New England Biolabs) on a QuantStudio 3 real-time PCR system (Thermo Fisher Scientific). The following primer pairs were used: Myc CAGGACTGTATGTGGAGCGG (forward) and GTCGTTGAGAGGGTAGGGGA (reverse); U6 CGCTTCGGCAGCACATATAC (forward) and AAAATATGGAACGCTTCACGA (reverse); 47S GCTGACACGCTGTCCTCTG(forward) and ACGCGCGAGAGAACAGCAG (reverse).

### Flow cytometry

HEK293T cells were stained with Live/Dead Fixable Near-IR Dead Cell Stain for 30 minutes at 4°C. Following staining, cells were washed once with PBS and centrifuged at 500g for 5 min. Cells were then fixed using the Foxp3 Transcription Factor Staining Buffer Kit for 1 h at room temperature. After fixation, cells were washed twice with permeabilization buffer and centrifuged at 1,200g for 5 min. The cells were incubated with anti-m^7^G antibody (1:1000) and anti-HA antibody (1:5000) at room temperature for 30 min. After two washes with permeabilization buffer and centrifugation at 1,200g for 5 minutes, cells were stained with the secondary antibodies at room temperature for 30 min. Cells were washed as previously described and analyzed by flow cytometry.

### *In Vitro* Methylation Assay

*In vitro* methylation assays were performed using SAM510™: SAM Methyltransferase Assay (G-Biosciences) according to the manufacturer’s instructions. Recombinant NSP14-NSP10 protein complex was incubated with GTP, ATP, UTP or CTP in the presence of S-adenosylmethionine at 37 °C. Reaction products were subsequently analyzed via LC-QToF to confirm the formation of m^7^GTP.

### Mass Spectrometry

100-200 ng of total RNA or de-capped poly(A) RNA from NSP14-transfected HEK293T cells were digested to nucleosides at 37°C overnight using a Nucleoside Digestion Mix (New England Biolabs). The digested RNAs were subsequently injected without prior purification on an Agilent 1290 Infinity II UHPLC equipped with a G7117 diode array detector and an Agilent 6495C Triple-Quadrupole Mass Spectrometer operating in positive electrospray ionization (+ESI) mode. UHPLC was conducted on a Waters XSelect HSS T3 XP column (2.1×100 mm, 2.5 µm) containing methanol and 10 mM ammonium acetate (pH 4.5) gradient mobile phase. Mass spectrometric data were acquired using dynamic multiple reaction monitoring (DMRM) mode. Each nucleoside species was identified based on the associated retention time and mass transition in the extracted chromatogram. Differentiations of isomeric nucleosides of methylated adenosine, guanosine and cytidine were described in previous publications^84,85^. For nucleoside quantification, calibration curves were constructed based on the mass responses of nucleoside standards with known concentrations, as determined by a UV spectrometer (Thermo Fisher Scientific, Evolution 220). Relative abundances were calculated by normalizing the amount of modified nucleoside to the corresponding unmodified nucleoside (e.g., m^7^G/G).

### RNA sequencing analysis

RNA-Seq libraries were prepared from poly(A)-enriched or m⁷G-MeRIP-immunoprecipitated RNA using NEBNext Ultra II RNA Library Prep Kit (New England Biolabs). Libraries were sequenced on an Illumina NovaSeq platform. Reads were aligned to the human genome (GRCh38) using STAR, and splicing events were quantified using rMATS and RSeQC. Intron retention and splice junction usage were compared across conditions.

### Alternative splicing, intron retention quantification and gene ontology analysis

Raw reads were adapter-trimmed with cutadapt v4.6 (minimum length 16 nt) to remove library-specific adapters. Splicing was quantified with VAST-TOOLS v2.5.1 against VastDB hg38 (Hs2). For alternative splicing (AS), PSI was computed per replicate, and ΔPSI was defined as the difference between condition means. For intron retention (RI), PIR/PSI was computed per replicate from VAST-TOOLS .IR2 files; replicates were retained if they met a VAST coverage rule (I5 ≥ 10 & I3 ≥ 5) or (I5 ≥ 5 & I3 ≥ 10), and EE ≥ 10, requiring ≥2/3 replicates with coverage per condition before computing condition means. Events were called differential when |ΔPSI| ≥ 15 and when replicate distributions showed no overlap ≥ 5 percentage points. Gene Ontology (GO) enrichment analysis was performed in R (v4.4.0) using clusterProfiler (v4.12.6). The input was the list of deduplicated genes with increased intron retention from VAST-TOOLS. Enrichment was run with enrichGO against org.Hs.eg.db (v3.19.1), using the full OrgDb annotation as background. Significance was defined as BH-adjusted p-value ≤ 0.05. "

### Computational evaluations

Computational analyses were based on the complex structures with PDB IDs 5C8S and 7QIF^86,87^. Models of NSP14 homologs were generated using Modeller, a widely used tool for homology modeling^88^. To perform ligand docking, GetCleft was first used to identify the binding cavity based on the original ligand positions. Subsequently, FlexAID was employed to dock the evaluated compounds into the defined cavity^89,90^. Both docking tools were used as implemented in NRGSuite-Qt^91^. To assess binding interactions, Surfaces was used to calculate per-residue contributions to the binding affinity between NSP14 and the original ligands GpppA and m^7^GpppG, as well as the docked compounds^92^.

### Statistical analyses

Data are presented as mean ± standard deviation (SD). Statistical comparisons between groups were made using a two-tailed unpaired Student’s t-test. A p-value < 0.05 was considered statistically significant. Statistical analyses were performed using GraphPad Prism 10.

## Acknowledgments

SARS-CoV-2 NSP10 and NSP14 constructs were a friendly gift from Adolfo García-Sastre at The Icahn School of Medicine at Mount Sinai, New York, NY. Compound 16 (#16) was a gift from Dr. Radim Nencka at the Czech Academy of Sciences in the Czech Republic. This work was supported by NIH Grant K22AI168257 (to J.C.-C.H), Rutgers Busch Biomedical Grant (to J.C.-C.H.) and RO1 AI166668 (to R.R.).

