## Supplemental Figures for "Coronavirus NSP14 Drives Internal m^7^G Modification to Rewire Host Splicing and Promote Viral Replication"

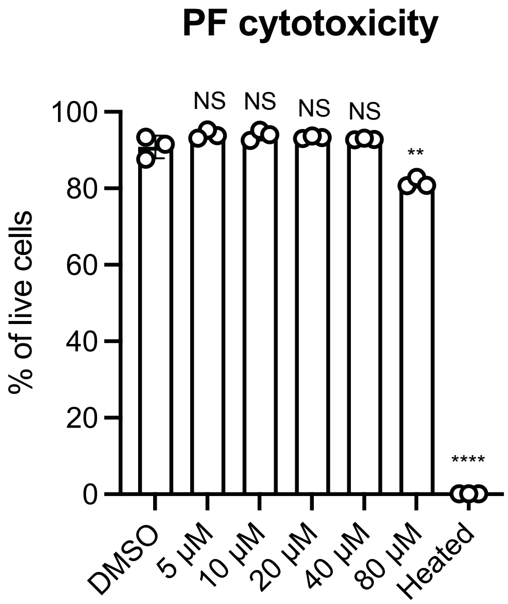


**Figure S1 Cytotoxicity assay of PF-03882845**
HEK293T cells were treated with increasing concentrations of PF-03882845 (PF) for 24 h. Cell viability was assessed using a cytotoxicity assay kit (Invitrogen; L34976) following the manufacturer's protocol and analyzed by flow cytometry. Data are presented as mean ± SD from three biological replicates. **P < 0.01, ****P < 0.001 by unpaired Student's t-test. NS, not significant.

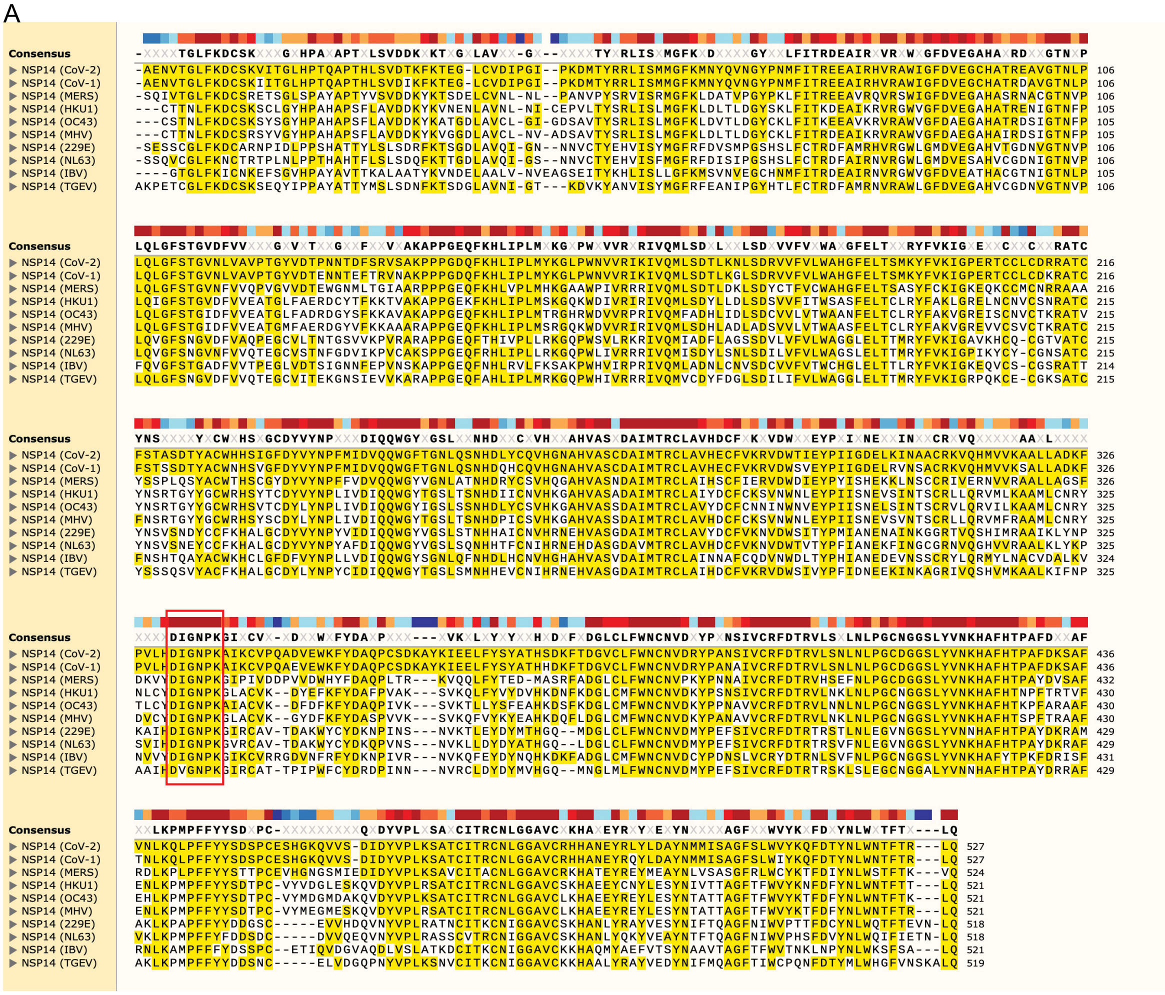


B

| Virus | Genus | Host | Protein seq |
|---|---|---|---|
| HCoV-229E | Alphacoronavirus | Human | NC_002645 |
| HCoV-NL63 | Alphacoronavirus | Human | NC_005831 |
| HCoV-HKU1 | Betacoronavirus | Human | NC_006577 |
| HCoV-OC43 | Betacoronavirus | Human | NC_006213 |
| MERS-CoV | Betacoronavirus | Human | NC_019843 |
| SARS-CoV-1 | Betacoronavirus | Human | NC_004718 |
| SARS-CoV-2 | Betacoronavirus | Human | NC_045512.2 |

C

| Similarity (%) | HCoV-229E | HCoV-NL63 | HCoV-HKU1 | HCoV-OC43 | MERS-CoV | SARS-CoV-1 | SARS-CoV-2 |
|---|---|---|---|---|---|---|---|
| HCoV-229E | X | | | | | | |
| HCoV-NL63 | 89 | X | | | | | |
| HCoV-HKU1 | 68 | 68 | X | | | | |
| HCoV-OC43 | 67 | 68 | 91 | X | | | |
| MERS-CoV | 66 | 68 | 74 | 74 | X | | |
| SARS-CoV-1 | 68 | 67 | 74 | 75 | 77 | X | |
| SARS-CoV-2 | 68 | 67 | 74 | 74 | 77 | 99 | X |

**Figure S2 Sequence analysis of coronavirus NSP14 proteins**

(A) Protein sequence alignment of NSP14 from representative coronaviruses. Conserved amino acid residues are highlighted in yellow, and the red box indicates the S-adenosylmethionine (SAM) binding motif. The alignment includes alphacoronaviruses (HCoV-229E [229E], HCoV-NL63 [NL63], transmissible gastroenteritis virus [TGEV]), betacoronaviruses (HCoV-HKU1 [HKU1], HCoV-OC43 [OC43], MERS-CoV [MERS], SARS-CoV [CoV-1], SARS-CoV-2 [CoV-2], mouse hepatitis virus [MHV]), and the gammacoronavirus infectious bronchitis virus (IBV).

(B-C) NSP14 sequence identity among human coronaviruses. Heatmap showing pairwise amino acid sequence similarity (%) of NSP14 across human coronaviruses.

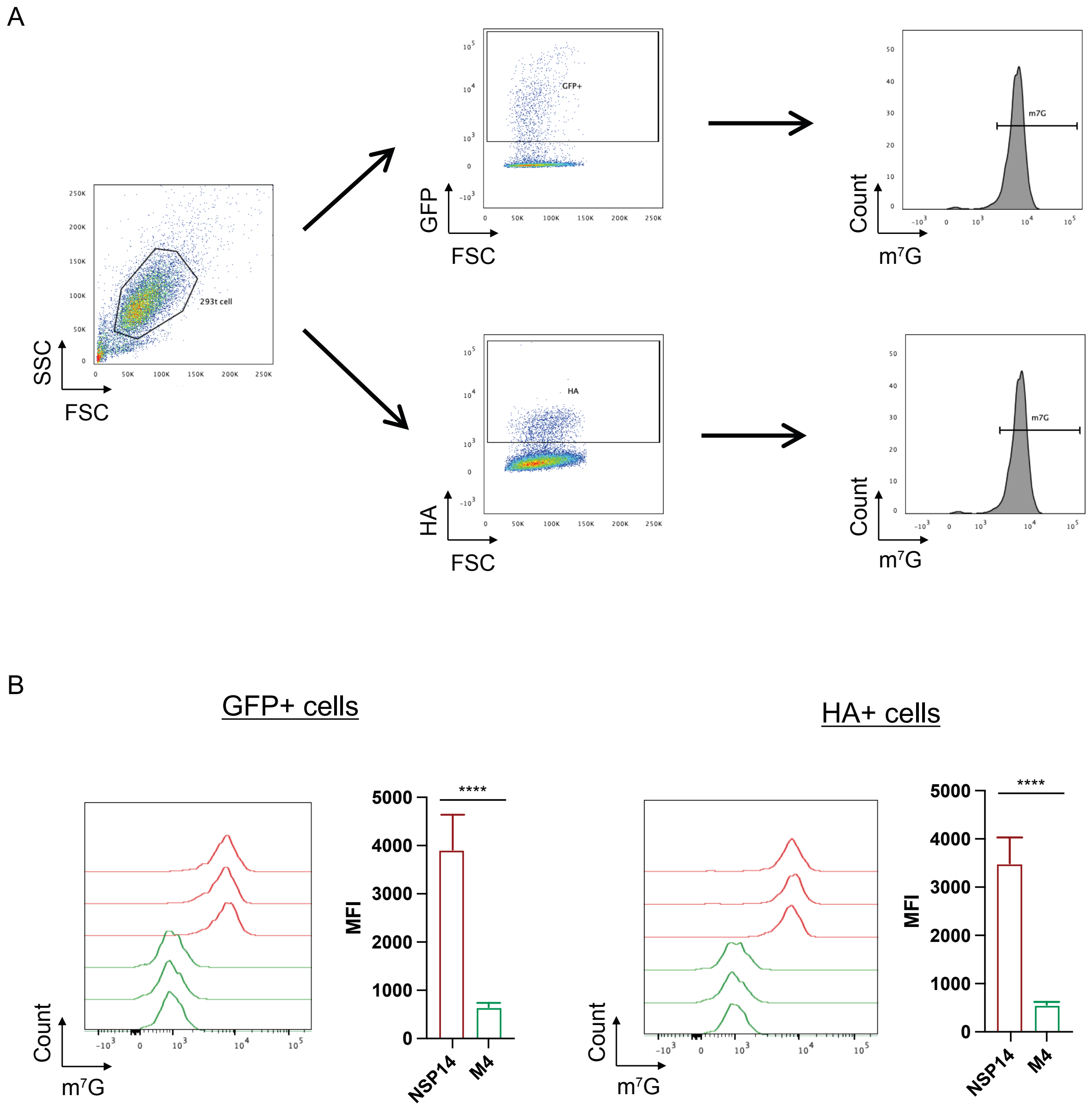


**Figure S3  Flow cytometry-based detection of RNA m$^7$G modification (m$^7$G FACS)**
(A) Schematic overview of the flow cytometry strategy used to detect m$^7$G modification. HEK293T cells were transfected with plasmids encoding HA- and GFP-tagged NSP14 for 24 h. Cells were fixed, permeabilized, and stained with antibodies against m$^7$G and HA, followed by flow cytometry analysis.

(B) Flow cytometry histograms and quantification of mean fluorescence intensity (MFI) showing m$^7$G signals in GFP$^+$ or HA$^+$ cells transfected with NSP14 or the *N7*-MTase mutant (M4). NSP14 significantly increased m$^7$G levels compared to the M4 mutant. Data are shown as mean ± SD from three biological replicates. ****$P < 0.0001$ by unpaired Student's t test.

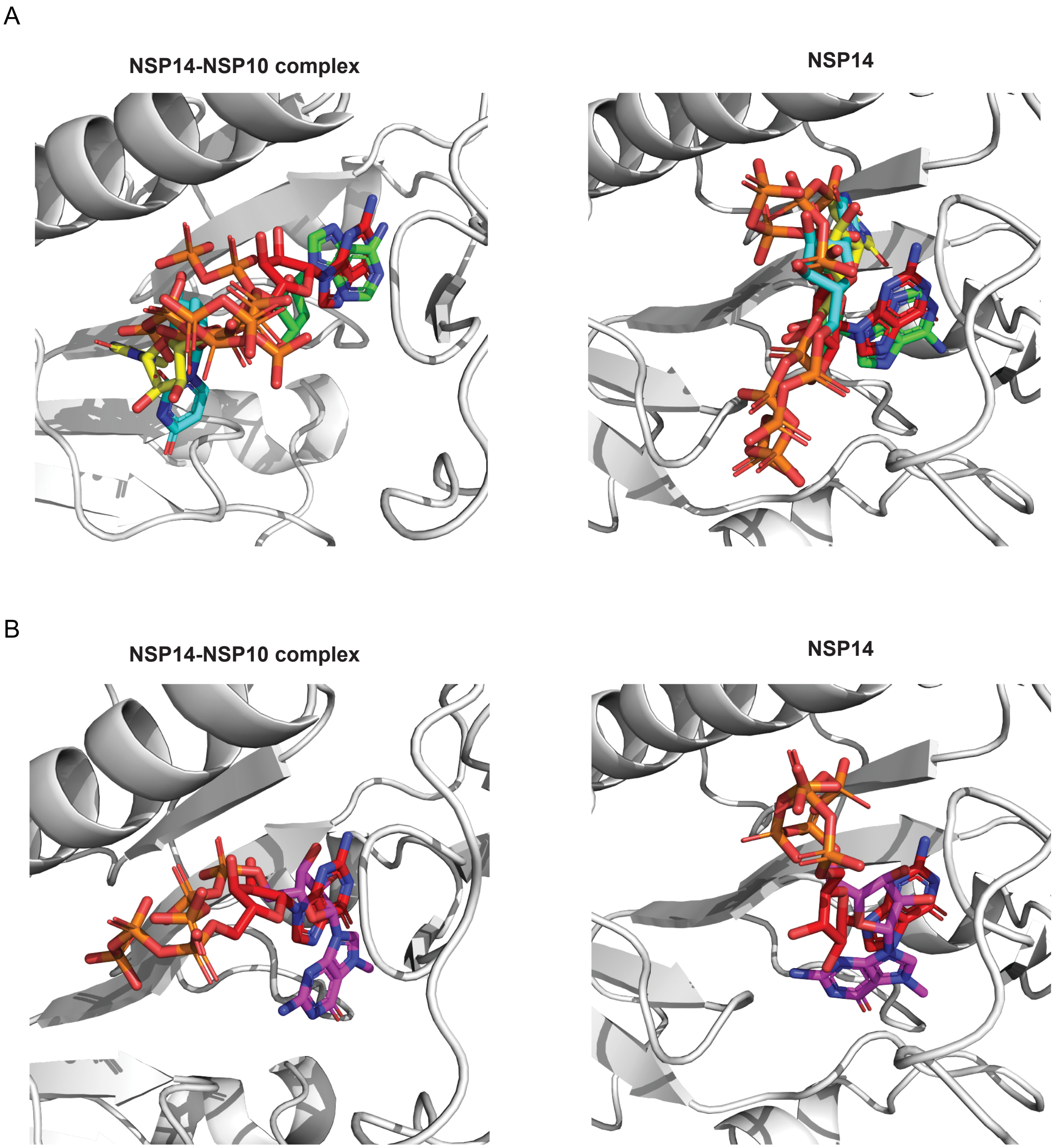


**Figure S4 Molecular docking analysis of nucleotide triphosphate binding to the NSP14-NSP10 complex and NSP14**

Structural modeling and docking were performed to assess the binding of nucleotide triphosphates to the NSP14-NSP10 complex and NSP14. Docking poses targeting a homology model based on the SARS-CoV-1 NSP14-NSP10 complex (PDB 5C8S) and the SARS-CoV-2 NSP14 structure bound to the cap analog $m^7G$ pppG (PDB 7QIF).

(A) Docking poses of GTP (red), ATP (green), UTP (cyan), and CTP (yellow) indicate that purine-based NTPs (GTP and ATP) align well within the binding cavity, while pyrimidine-based NTPs (UTP and CTP) are displaced, suggesting reduced binding compatibility.

(B) Comparison of docking poses of GTP (red) and $m^7$GTP (magenta) reveals that the *N*7-methylated guanine of $m^7$GTP extends outside the binding cavity, consistent with its role as a reaction product rather than a substrate.

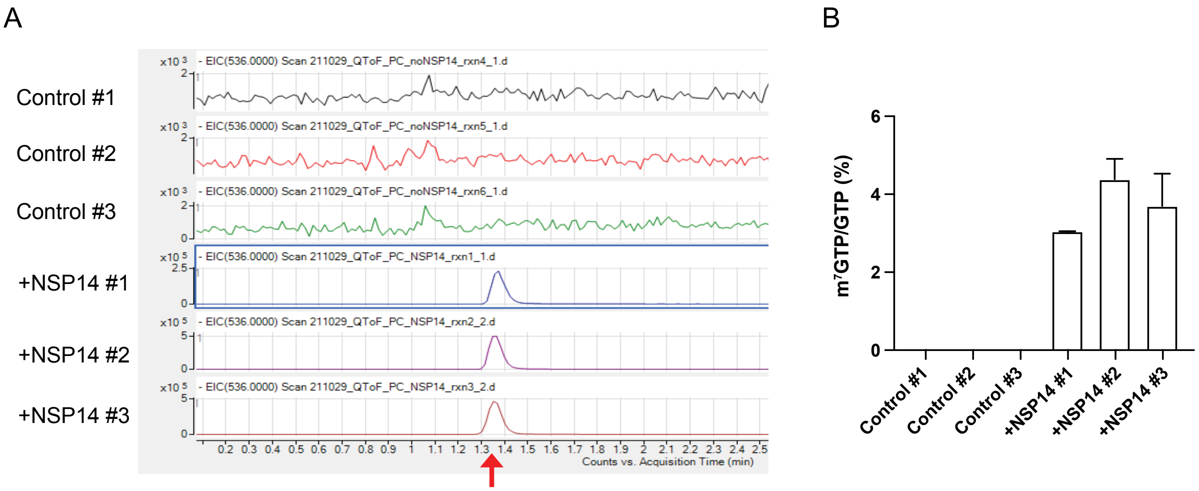


**Figure S5 Mass spectrometry confirms NSP14-mediated synthesis of m$^7$GTP**

(A) Extracted ion chromatograms from liquid chromatography-quadrupole time-of-flight (LC-QToF) mass spectrometry showing detection of m$^7$GTP in *in vitro* methylation reactions with or without recombinant NSP14-NSP10 complex. A distinct peak corresponding to m$^7$GTP (red arrow) was detected in all NSP14-treated samples but absent in control reactions.

(B) Quantification of m$^7$GTP as a percentage of total GTP from three biological replicates. Data are presented as mean ± SD.

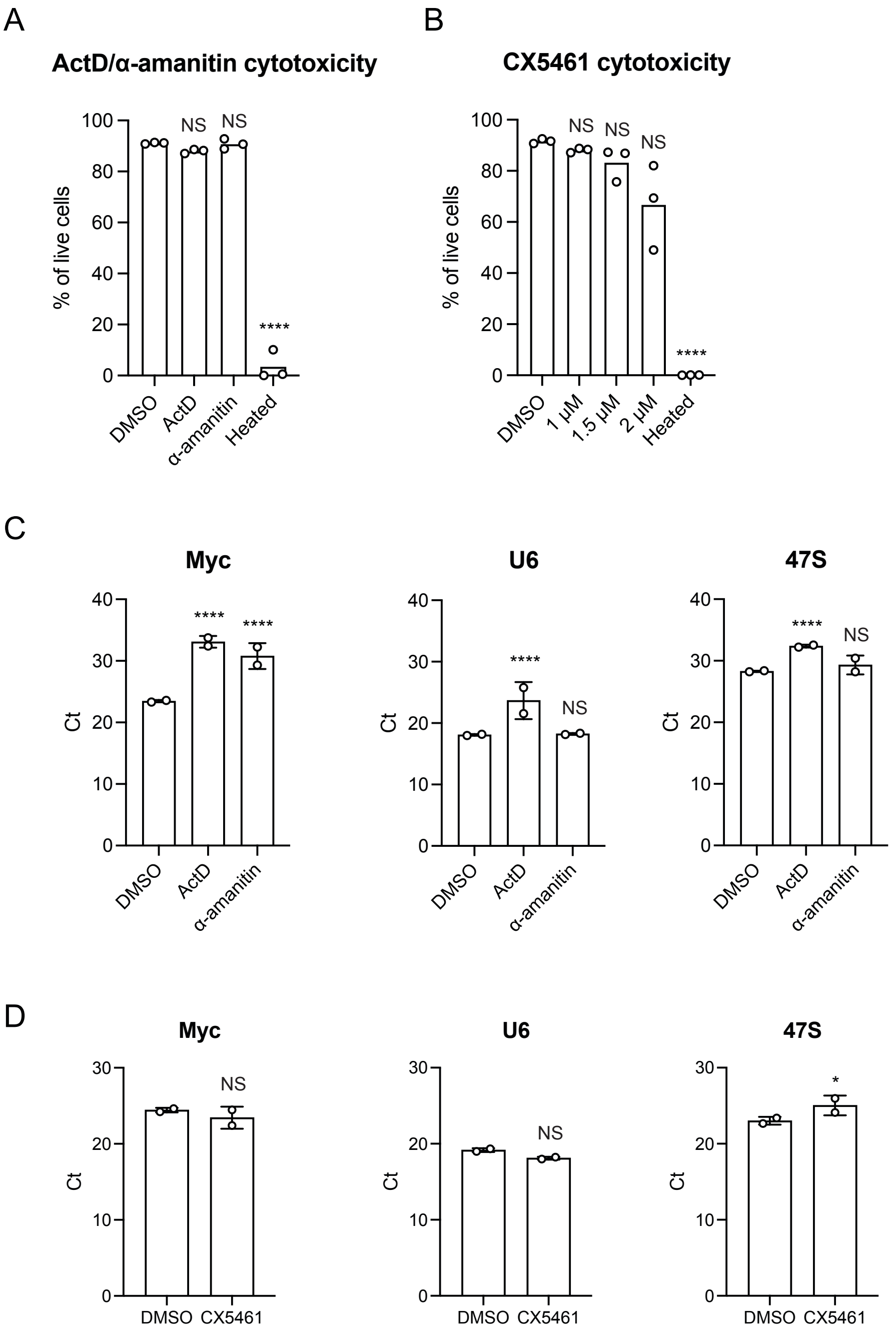


**Figure S6 Cytotoxicity assay of transcription inhibitors**

HEK293T cells were treated with actinomycin D, α-amanitin (A and C) and CX-5461 (B and D) for 24 h. (A, B) Cell viability was assessed using a cytotoxicity assay kit (Invitrogen; L34976) following the manufacturer's protocol and analyzed by flow cytometry. (C, D) RNA levels of indicated genes in HEK293 T cells treated with transcription inhibitors was determined by RT-qPCR. Data are presented as mean ± SD from two to three biological replicates. *$P < 0.05$, ****$P < 0.001$ by unpaired Student's t-test. NS, not significant.

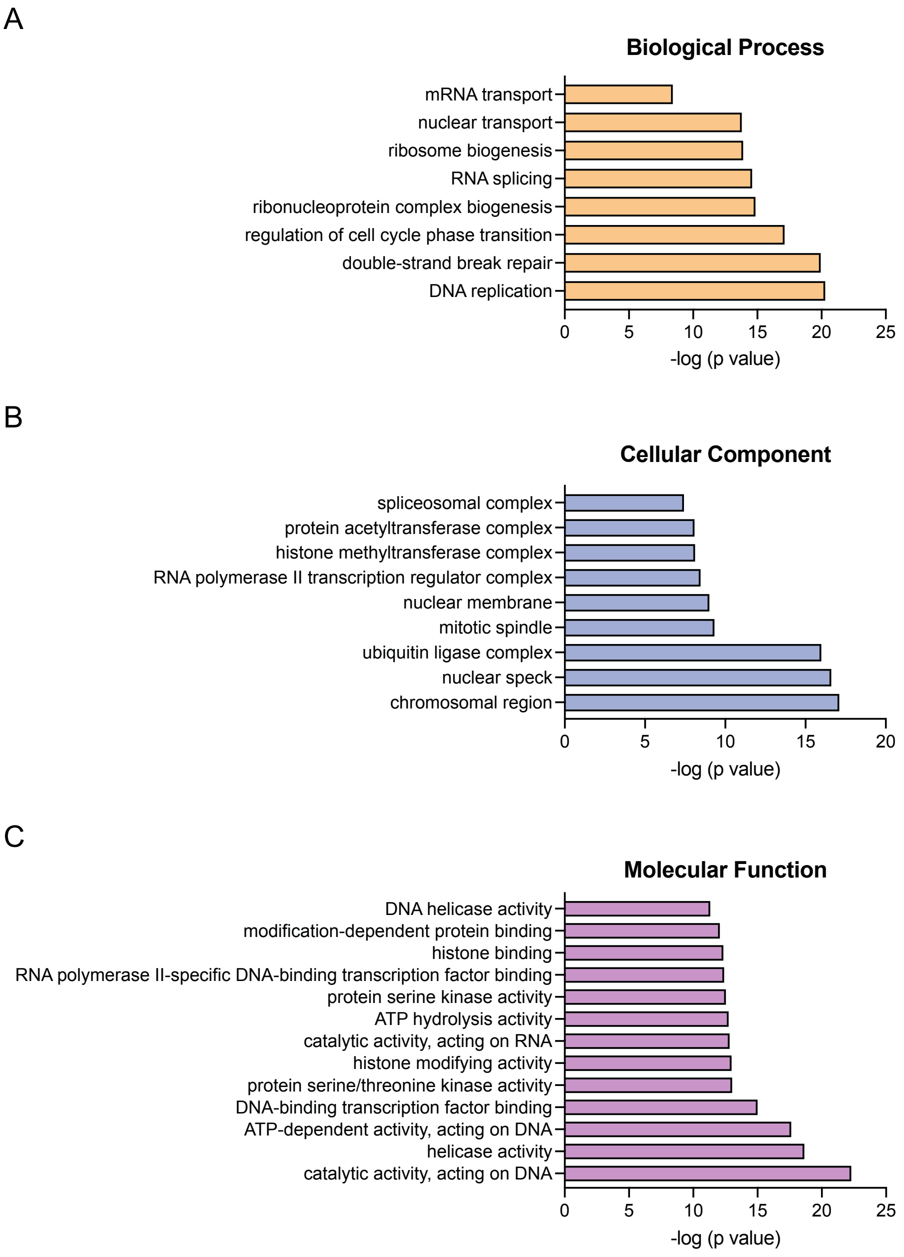


**Figure S7 Gene ontology (GO) enrichment analysis of genes with increased intron retention upon NSP14 expression.**

(A) Biological Process terms show enrichment for pathways involved in genome maintenance and RNA metabolism, including DNA replication/repair, cell cycle regulation and RNA processing. (B) Cellular Component terms highlight enrichment in nuclear structures such as chromosomal regions, spliceosomal complexes and nuclear specks, consistent with roles in chromatin organization and RNA processing. (C) Molecular Function terms are enriched for DNA- and RNA-related enzymatic activities, helicases, transcription factor binding, and chromatin-modifying enzymes. Together, these results indicate that NSP14 disrupts host pathways coordinating genome stability, RNA splicing and transcriptional regulation.

A

| Gene | Coordination | Intron size (bp) | Exons | dPSI | IR | RT-PCR amplicon size | |
|---|---|---|---|---|---|---|---|
| | | | | | | Long (bp) | Short (bp) |
| PRRC2A | chr6:31637325-31637445 | 121 | 30 and 31 | 36.54 | UP | 204 | 83 |
| ATG101 | chr12:52070243-52070353 | 111 | 1 and 2 | 35.84 | UP | 201 | 90 |
| RPS27A | chr2:55232709-55232807 | 99 | 1 and 2 | 29.78 | UP | 185 | 86 |
| MAFG | chr17:81923058-81923149 | 92 | 2 and 3 | 37.92 | UP | 200 | 108 |
| SSBP3 | chr1:54228357-54228448 | 92 | 16 and 17 | -0.44 | Control | 199 | 107 |

B

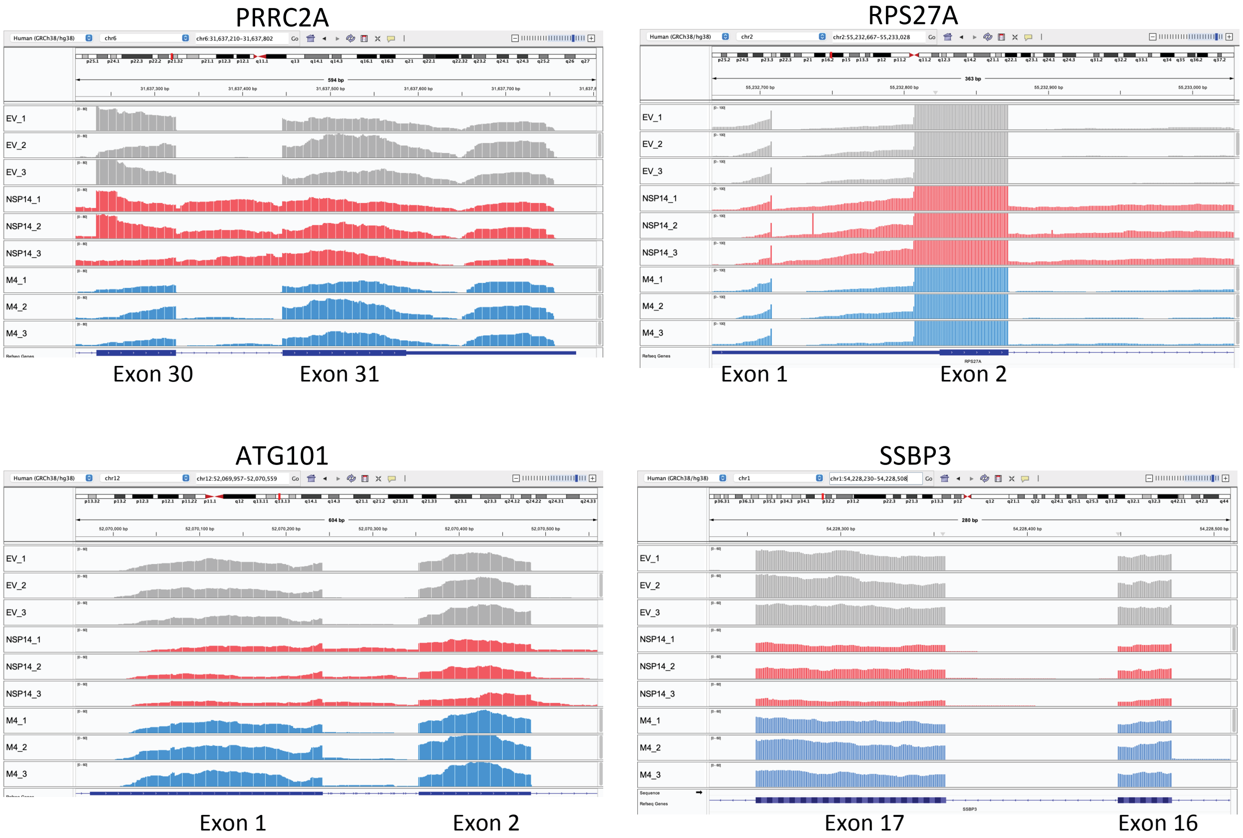


**Figure S8 RNA-Seq analysis of NSP14-induced intron retention**
(A) Selected genes analyzed for intron retention, including genomic coordinates, intron size, involved exons, difference in percent spliced-in index (dPSI), intron retention (IR) status, and RT-PCR amplicon sizes. PRRC2A, ATG101, and RPS27A displayed increased intron retention upon NSP14 expression, while SSBP3 served as a control with no significant changes. Data extracted from Zaffagni *et al.*, 2022.

(B) Representative IGV alignment tracks of PRRC2A, RPS27A, ATG101, and SSBP3 showing intron retention. RNA-seq read coverage is displayed for cells transfected with empty vector (EV, gray), wild-type NSP14 (red), and the M4 mutant (blue). Increased intron retention is observed in PRRC2A, RPS27A, and ATG101 upon NSP14 expression, while SSBP3 shows no change and serves as a negative control.

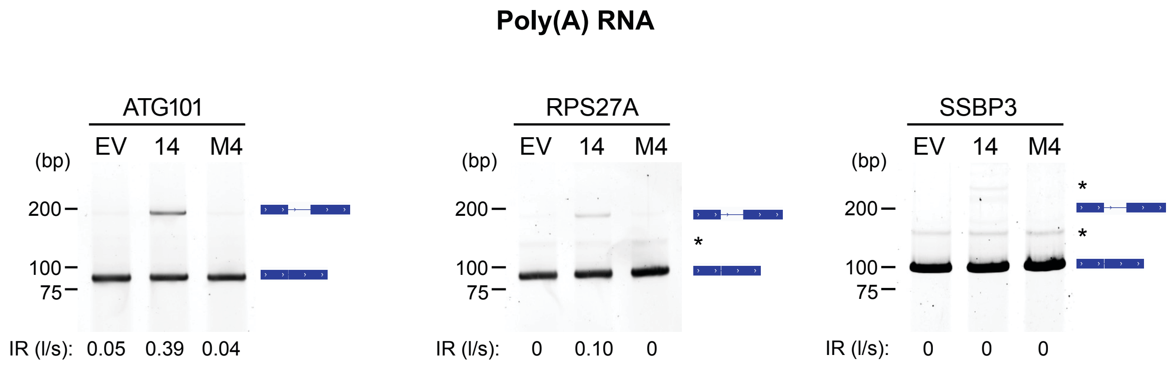


**Figure S9 RT-PCR analysis of intron retention in poly(A) RNA**
HEK293T cells were transfected with EV, NSP14, or M4 for 24 h. Poly(A) RNA was reverse-transcribed and amplified by PCR. The resulting PCR products were analyzed by DNA gel electrophoresis. The intron retention ratio of unspliced “long” fragments to spliced “short” fragments (IR(l/s)) was quantified using ImageJ. *, non-specific PCR product.

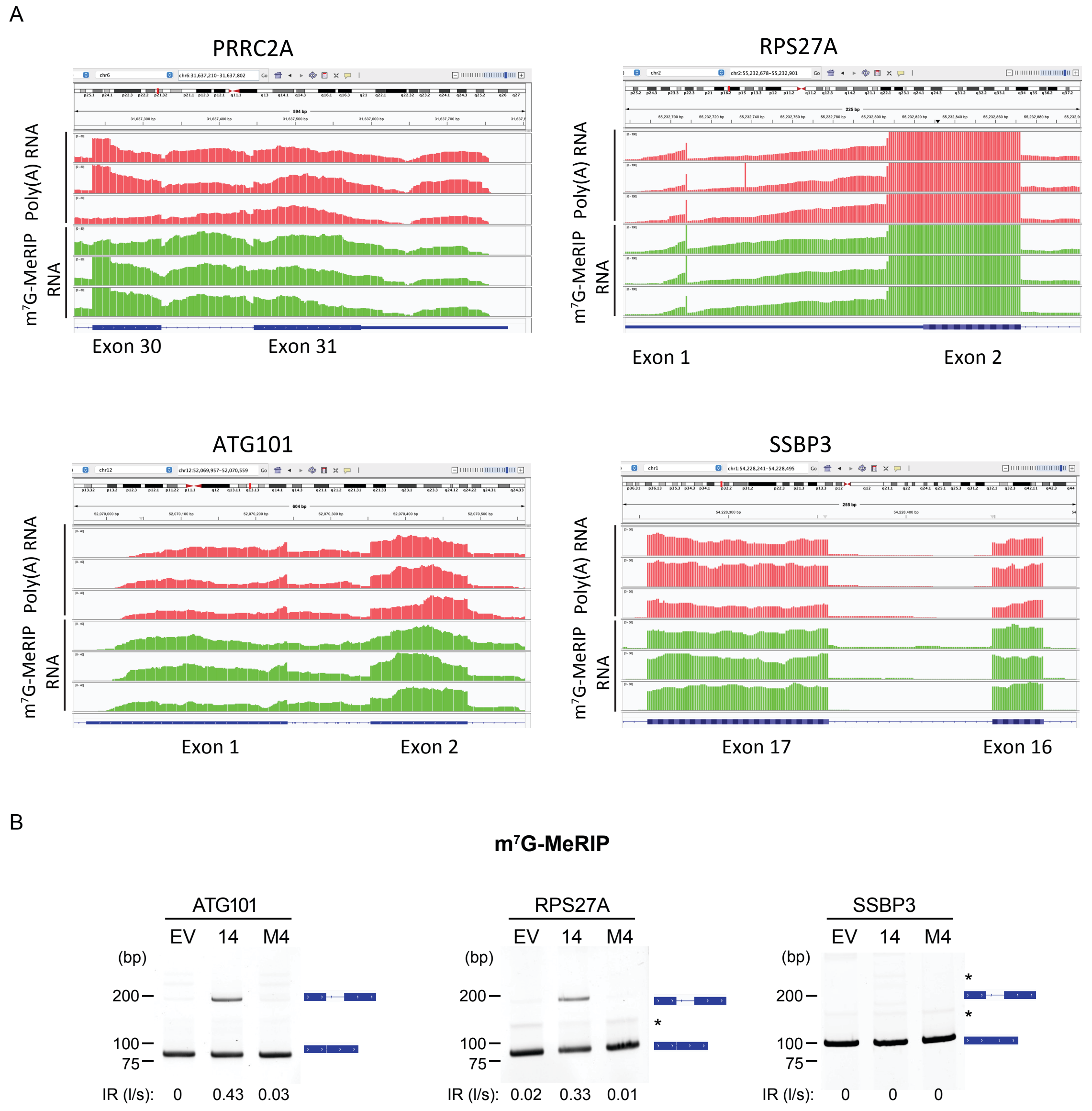


**Figure S10 NSP14-induced intron retention is enriched in m$^7$G-modified RNA**
(A) Representative IGV alignment tracks of PRRC2A, RPS27A, ATG101, and SSBP3, comparing m$^7$G-MeRIP RNA and poly(A) RNA. Read coverage is shown for cells transfected with empty vector (EV, gray), NSP14 (red), and the M4 mutant (blue). Notably, increased intron retention is observed in the m$^7$G-MeRIP RNA compared to the poly(A) RNA.

(B) Validation of NSP14-induced intron retention by RT-PCR from m$^7$G-MeRIP RNA. HEK293T cells were transfected with EV, NSP14, or M4 mutant for 24 h. Purified m$^7$G-modified RNA was reverse-transcribed and amplified with primers flanking retained introns. The resulting PCR products were analyzed by DNA gel electrophoresis. The intron retention ratio of unspliced “long” fragments to spliced “short” fragments (IR(l/s)) was quantified using ImageJ.  *, non-specific PCR product.

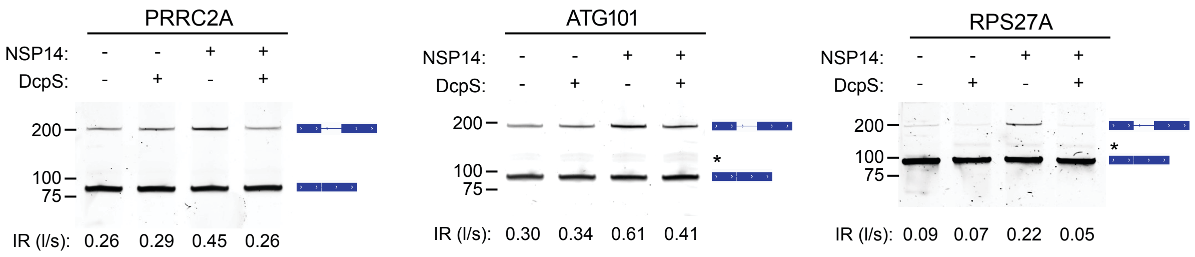


**Figure S11  RT-PCR analysis of intron retention in total RNA**

HEK293T cells were transfected with EV, NSP14, or M4 for 24 h. Total RNA was reverse-transcribed and amplified by PCR. The resulting PCR products were analyzed by DNA gel electrophoresis. The intron retention ratio of unspliced "long" fragments to spliced "short" fragments (IR(I/s)) was quantified using ImageJ. *, non-specific PCR product.

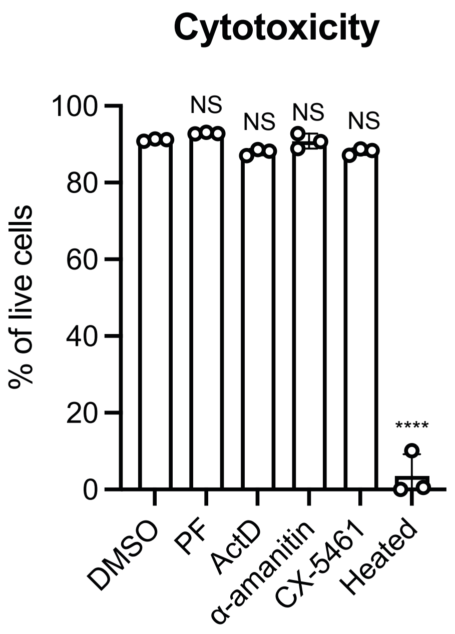


**Figure S12 Cytotoxicity assay of inhibitors**
Vero E6 cells were treated with PF-03882845 (PF) and transcription inhibitors (actinomycin D [ActD], α-amanitin, CX-5461) for 24 h. Cell viability was assessed using a cytotoxicity assay kit (Invitrogen; L34976) following the manufacturer's protocol and analyzed by flow cytometry. Data are presented as mean ± SD from three biological replicates. ****$P < 0.001$ by unpaired Student's t-test. NS, not significant.
